# Macrophage mediated mesoscale brain mechanical homeostasis mechanically imaged via optical tweezers and Brillouin microscopy *in vivo*

**DOI:** 10.1101/2023.12.27.573380

**Authors:** Woong Young So, Bailey Johnson, Patricia B. Gordon, Kevin S. Bishop, Hyeyeon Gong, Hannah A Burr, Jack Rory Staunton, Chenchen Handler, Raman Sood, Giuliano Scarcelli, Kandice Tanner

## Abstract

Tissues are active materials where epithelial turnover, immune surveillance, and remodeling of stromal cells such as macrophages all regulate form and function. Scattering modalities such as Brillouin microscopy (BM) can non-invasively access mechanical signatures at GHz. However, our traditional understanding of tissue material properties is derived mainly from modalities which probe mechanical properties at different frequencies. Thus, reconciling measurements amongst these modalities remains an active area. Here, we compare optical tweezer active microrheology (OT-AMR) and Brillouin microscopy (BM) to longitudinally map brain development in the larval zebrafish. We determine that each measurement is able to detect a mechanical signature linked to functional units of the brain. We demonstrate that the corrected BM-Longitudinal modulus using a density factor correlates well with OT-AMR storage modulus at lower frequencies. We also show that the brain tissue mechanical properties are dependent on both the neuronal architecture and the presence of macrophages. Moreover, the BM technique is able to delineate the contributions to mechanical properties of the macrophage from that due to colony stimulating factor 1 receptor (CSF1R) mediated stromal remodeling. Here, our data suggest that macrophage remodeling is instrumental in the maintenance of tissue mechanical homeostasis during development. Moreover, the strong agreement between the OT-AM and BM further demonstrates that scattering-based technique is sensitive to both large and minute structural modification *in vivo*.

## Main

Organs have distinct mechanical properties[1, 2]. For example, the Young’s modulus of the brain is ∼200Pa whereas comparable values are ∼ 10kPa for muscle tissue[1–3]. Within a given organ, there are regional differences in mechanical properties that are linked to functional properties[3, 4]. A mechanical readout in one functional unit may correspond to normal physiology whereas a comparable measurement in a different unit may indicate the presence of disease. Methods to measure tissue mechanics largely rely on techniques that quantitate the elastic response usually a Young’s modulus averaged over large regions[3]. Moreover, mechanical properties are probed at different temporal and force scales[5]. One area of active exploration focuses on what do measurements of “compliance” or “stiffness”, “solid-like” or “liquid-like” mean when considering a cell or tissues[5, 6]. Simply, can we link a quantitative mechanical measurement to an underlying biological process? A stand-alone method to assess what aspects of tissue mechanics are relevant for diagnostics or biological mechanisms remains an ongoing process.

Organ homeostasis requires an intricate balance between diverse and dynamic tissue constituents[7, 8]. Within the brain, homeostasis is possible due to specialized networks of diverse neuronal architectures, protected by specialized vasculature systems such as the blood-brain barrier and meningeal lymphatics and shaped by tissue resident immune cells[9–12]. During development, tissue-resident immune cells such as microglia play a significant role in sculpting the developing brain and neuronal connectivity to establish and maintain normal brain function[13]. Not restricted to developmental processes, these stromal cells also impact the onset of different pathologies such as cancers and neurodegenerative diseases [14–17]. As first suggested by D’Arcy Thompson more than a century ago, biophysical properties of tissue are also key determinants of development[18]. However, probing these in large intact tissues with sub-cellular resolution non-invasively remains challenging. There are several techniques that allow measurement of shear, young’s or longitudinal moduli as a function of different length, frequency and force scales. But tissue anisotropy complicates our ability to reliably move interchangeably between these different methods. Scattering based techniques such as Brillouin microscopy have long been used to probe biological phenomena[19–28]. However, our conventional understanding of materials is determined with tools like optical tweezers and atomic force microscopy at lower frequencies. Thus, connecting measurements from these established methods to measured material properties under gigahertz frequencies is needed to understand the full frequency response for viscoelastic materials. Here, we compared two independent mechanical mapping methods to compare/contrast the measured physical property and the underlying physical structure. Here, we focused our studies on optically probing brain morphogenesis to decipher the interplay between a mechanical signature and stromal dynamics.

The zebrafish has long been utilized as a model organism as many organs are conserved in mammalian counterparts. Moreover, the transparency of larval zebrafish allows for non-invasive optical techniques[22, 29–35]. Here, we applied two optical based techniques: optical tweezer based active microrheology (OT-AMR) and Brillouin microscopy in an animal model that recapitulates spatial anisotropy and temporal evolution of mechanical properties in native intact tissues (Figure 1a, b). With these data, we compared the microscale shear and longitudinal moduli for each of the modalities to map development for the same animal over days. We also calculated the relative density factors to convert the measure Brillouin shifts into longitudinal moduli. Our data revealed a strong correlation between the Brillouin-longitudinal modulus and the elastic moduli measured at lower frequencies. We then aimed to determine the correlation between functional units of the brain with the observed mechanical signatures.

**Figure 1:**
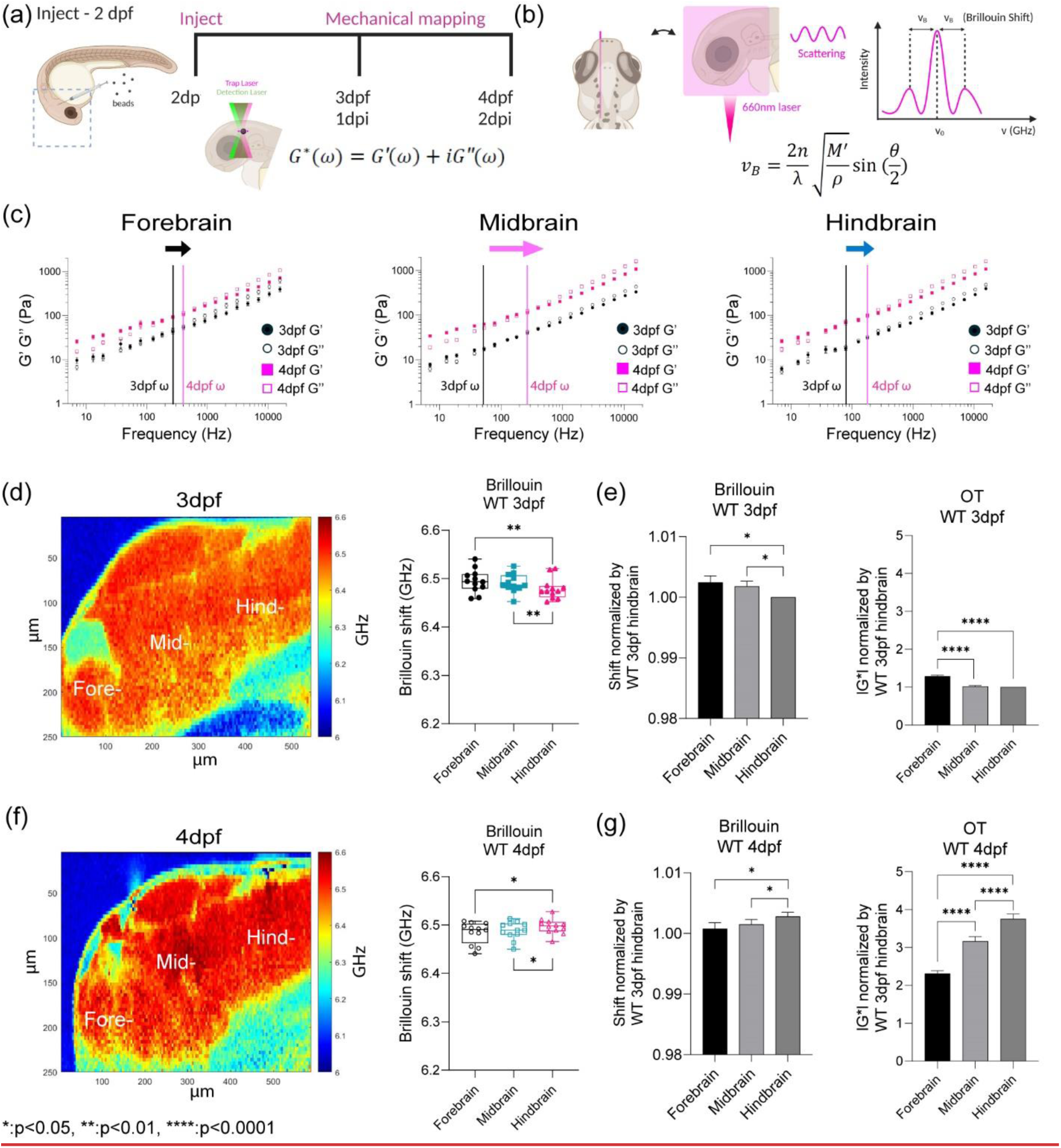
Optical trap active microrheology and Brillouin microscopy show strong correlation for measured microscale mechanics of functionally distinct units of the brain. (a) Schematic of brain mechanics measurement for optical trap where 1 µm beads are directly injected into midbrain parenchyma for mechanical mapping at 3 days-post-fertilization (3dpf) / 1 day post injection (dpi) and 4dpf / 2dpi. High power trap laser oscillates the bead while the stationary low power detection laser records the bead motion, which is translated into complex modulus (G*), elastic modulus (G’), and viscous modulus (G’’) (b) Schematic of Brillouin microscopy measurement on brain parenchyma where the scattering is collected upon 660nm laser radiation (c) log-log plot of WT brain mechanics (elastic modulus, G’, and viscous modulus, G’’) and frequencies (7Hz to 15kHz) in terms of development (3dpf and 4dpf) and brain regions (forebrain, midbrain, and hindbrain) with crossover frequency labelled where viscous modulus becomes greater than elastic modulus; 3dpf in black and 4dpf in magenta. Error bars in standard of error (3dpf = 4 fish - forebrain n (number of beads/fish) = 78, midbrain n = 90, and hindbrain n= 77. 4dpf = 6 fish – forebrain n (number of beads/fish) = 202, midbrain n = 224, and hindbrain n= 259) (d) 3dpf WT Brillouin microscopy image with Brillouin shift (GHz) in terms of ‘Fore-‘, ‘Mid-‘, and ‘Hind-‘ (n = 11). ** p<0.01, paired two-tailed t-tests. (e) Normalized bar graph of Brillouin microscopy on WT 3dpf in respect to average Brillouin shift (GHz) of WT 3dpf hindbrain and normalized bar graph of WT 3dpf complex modulus (G*) from optical tweezer in respect to WT 3dpf hindbrain based on 19 different frequencies. * p<0.05, and **** p<0.0001, paired two-tailed t-tests. (f) 4dpf WT Brillouin microscopy image with Brillouin shift in terms of ‘Fore-‘, ‘Mid-‘, and ‘Hind-‘ (n = 12). * p<0.05, paired two-tailed t-tests. (g) Normalized bar graph of Brillouin microscopy on WT 4dpf in respect to average Brillouin shift (GHz) of WT 3dpf hindbrain and normalized bar graph of WT 4dpf complex modulus (G*) from optical tweezer in respect to WT 3dpf hindbrain based on 19 different frequencies. * p<0.05, and **** p<0.0001, paired two-tailed t-tests.

Using a suite of tools where we either chemo-genetically or chemically ablated stromal constituents, we probed the role of mechanical mediated tissue remodeling in larval zebrafish. Tissue viscoelasticity was driven by both macrophages and csf1r receptor mediated crosstalk within the developing brain. Our data provide evidence to link viscoelastic properties of tissue to macrophage mediated tensional homeostasis in the brain. These data also suggest that stromal dynamics that induce acute changes in mechanical properties may play an augmented role in changes in both normal tissue architecture and for therapies targeted against them.

### Optical trap active microrheology and Brillouin microscopy show strong correlation for measured microscale mechanics of functionally distinct units of the brain

The developmental program that drives the formation of the architecturally distinct forebrain, midbrain and hindbrain and blood brain barrier are conserved between zebrafish and mammals, and it is heavily reliant on the balance between mechanical forces and tissue stiffness[35, 36] (Extended Data 1a). To measure the mechanical properties of the developing brain from 3 - 4dpf, we employed optical trap based active microrheology (OT-AMR) (Figure 1a, c). First, we introduced 1 micron-diameter polystyrene spheres directly into the brain parenchyma at 2 days post fertilization (2dpf). Using these spheres as mechanical probes, we applied a multiplexed-sinusoidal oscillation based on ∼nm amplitude to each bead over a range of frequencies from 7Hz to 15kHz. The displacements of the bead are simultaneously tracked via back focal plane interferometry. Quantitation of the local viscoelasticity, G*(ω) at that bead position is achieved by correlating the trap and bead positions. The complex modulus, G*(ω) = |G*|exp(iδ) = G′(ω) +*i*G″(ω) which then gives storage (G’(ω) and loss moduli (G”(ω). Then, comparing G’ (elastic modulus) and G’’ (viscous modulus) by G’’/G’ would result into loss tangent or hystersivity to determine the relative state between solid-like (more G’ contribution) or liquid-like characteristics (more G’’ contribution) based on crossover frequency. We show that brain tissue is mechanically heterogeneous as a function of region of the brain and age (Figure 1c, Extended Data 1b). At 3 dpf, the midbrain is more liquid-like than the hind and forebrain respectively. In contrast, the forebrain is more rigid than the mid- and hindbrain. As the brain develops, all regions become more solid-like with the greatest transition occurring for the mid brain. An increase in the complex modulus is observed for all regions with the hind brain showing the greatest increase. Principles modeled on non-Newtonian fluids have been used to model rheological properties of tissue. One such concept is that viscoelasticity (complex moduli (G*)) obeys frequency-dependent power laws, |G*(ω)| = aω^b^, where the dependence b varies for different frequency regimes and different cell types[37–39]. Here, G’ and G” monotonically increase as a function of frequency and show power law dependence at frequencies >400Hz (Figure 1c, Extended Data 1b and c). Using the metrics, regional comparison at 3dpf showed that the slope of the line is greatest for the hindbrain (0.600) followed by the forebrain(0.582) and then the midbrain (0.565) where each of the values fall between 0.5 and 0.75 which correspond to formulaic descriptions of flexible and semi-flexible polymers respectively. As the animal ages, the mid-brain shows the greatest change where the slope of the line now changes from 0.565 to 0.607. The normalized complex modulus data from whole frequency range by 3 dpf hindbrain shows that each brain regions has distinct mechanical property with statistical significance based on paired t-test (Figure 1e and g, Extended Data 1d and e). All regions stiffen as a function of age but not uniformly where the hindbrain shows the greatest increase in stiffness.

OT-AMR offers the advantage of quantifying absolute values of viscoelasticity[29, 40]. However, the technique necessitates the introduction of probes, that if introduced in sufficient numbers can perturb normal physiology. Mechanical phenotypes of large tissues can only be obtained by averaging across many samples assuming random bead distribution. Thus, assessing the viscoelasticity of an individual fish is difficult. Brillouin microscopy (BM) can overcome this challenge as it is a purely non-invasive technique, based on the intrinsic inelastic scattering due to incident photon-phonon interactions in materials, whose spectral characterization provides information correlated with the underlying mechanical properties (Figure 1b)[25, 28]. Formulaically, the frequency shift (*ν*_*B*_) of Brillouin scattered photons (on the order of GHz) relate to the local refractive index (n), mass density (*ρ*), and the longitudinal elastic modulus (M’) of samples (Figure 1b)[25, 28]. Thus, stiffer samples would result in higher frequency shifts. We determined that Brillouin shifts were comparable in detecting mechanical heterogeneity of brain regions at 3 and 4dpf (Figure 1d and f). Furthermore, the trends in stiffness at 3 and 4 dpf based on Brillouin microscopy are the same as those measured using OT-AMR (Figure 1d-g). At 3 dpf, hindbrain is the softest but becomes the stiffest at 4dpf with a significant increase in frequency shift (Figure 1e and g, Extended Data 1f).

### Brillouin longitudinal modulus using calculated density factor shows greater correlation for OT measurements obtained at lower frequencies in the zebrafish brain

Mechanical heterogeneity is comparable at different temporal scales for BM (∼GHz) and OT-AMR (7Hz – 15kHz) *in vivo*. We then aimed to link tissue architecture that provide the contrast for the scattering signals measured by BM (Figure 2a). One of the most dramatic changes is that the neuronal networks increase in size and complexity as the brain develops. Recently, it has been proposed that the acetylation of lysin 40 of α-tubulin, a key component of neurons, is important for stiffness of developing tissue as well as cell mechanics [41, 42]. Whole-mount immunofluorescence of acetylated tubulin of the brain shows differential structure and clustering of neuronal networks between 3 and 4 dpf, especially in the midbrain and hindbrain. An increase in normalized intensity of acetylated tubulin was observed for all brain regions during development (Figure 2b-d). BM has been recently used to non-invasively infer the mechanical properties of engineered thick tissues such as organoids and in living animals[25, 26, 28]. Converting the measured Brillouin shift into a longitudinal modulus requires that additional optical and material properties are known [20]. Thus, we next quantified the relative areas of the different regions of the brain as well as the number of cells to determine a correction of the density factor for all of the cells present (Figure 2e-g). During development, not only are resident cells increasing in numbers but there are infiltrating cells such as macrophages (Figure 2g). We next estimated the differences in these infiltrating cells by quantifying macrophages at the mid-brain as a function of development (Figure 2g). Moreover, there are also differences in the timescale between the OT measurements and that due to the Brillouin phenomenon. Therefore, we compared the raw and corrected correlation between Brillouin (longitudinal) modulus, M’ and OT at 7Hz, 907 Hz and 15kHz in terms of the elastic modulus G’, for development stages during 3-4dpf (Figure 2h-i, Extended data 2). Comparison of linear fits between log(G’) for forebrain, midbrain, and hindbrain) and log(calculated Brillouin modulus M’) revealed poor correlation between techniques before corrections based on estimated brain refractive index and density (Figure 2h and i, Extended data 2). While the overall correlations improved post correction, better fits were obtained for values measured at 7 and 907Hz compared to 15kHz (Figure 2h and i, Extended data 2).

**Figure 2:**
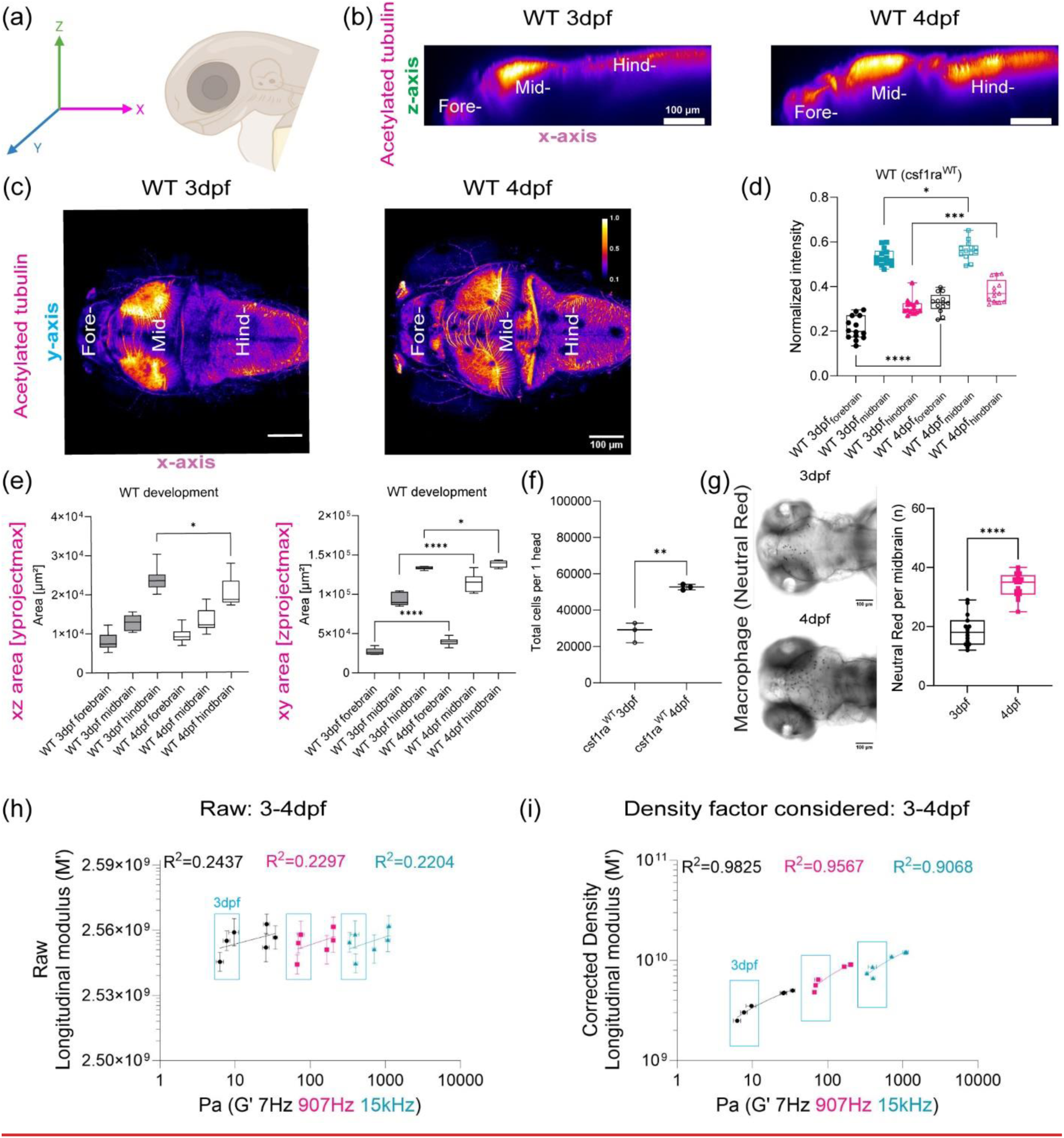
Brillouin longitudinal modulus using calculated density factor shows greater correlation for OT measurements obtained at lower frequencies *in vivo*. (a) Schematic of immunofluorescence of acetylated tubulin in different axis projections, z-axis and y-axis focusing on brain parenchyma excluding eye (b) y-axis maximum projection of WT acetylated tubulin over the development (c) z-axis maximum projection of WT acetylated tubulin intensity excluding eye over the development (d) Quantitation of normalized acetylated tubulin intensity in terms of development and brain regions (WT 3dpf: n = 14, WT 4dpf: n = 13). * p<0.05, *** p<0.001, **** p<0.0001, unpaired two-tailed t-tests. (e) Quantitation of brain region area based on acetylated tubulin in terms of development, xz (y-axis maximum projection), and xy (z-axis maximum projection) (WT 3dpf: n = 14, WT 4dpf: n = 13). * p<0.05, **** p<0.0001, unpaired two-tailed t-tests. (f) Dissociated head cells per fish based on the cluster of 15 fish in terms of development (WT 3dpf: n = 45, WT 4dpf: n = 45). Each dot represents a group of 15 fish head. ** p<0.01, unpaired two-tailed t-tests. (g) Macrophages at midbrain labelled by neutral red assay for WT in terms of development (WT 3dpf: n = 15 and WT 4dpf: n = 18). **** p<0.0001, unpaired two-tailed t-tests. (h) Raw correlation between corrected Brillouin (longitudinal) modulus (M’) and OT elastic modulus (G’) at 7Hz, 907Hz, and 15kHz during development (3-4dpf) in log-log plot (i) Density factor corrected correlation between corrected Brillouin (longitudinal) modulus (M’) and OT elastic modulus (G’) at 7Hz, 907Hz, and 15kHz during development (3-4dpf) in log-log plot.

### OT and Brillouin microscopy can detect changes in tissue mechanical properties in CSF1r zebrafish mutant with delayed macrophage invasion

As we observed difference in macrophage numbers as a function of development, we then asked if the presence of these cells also contributed to the overall mechanical properties of the brain. Moreover, if OT and Brillouin would be sensitive to microscale perturbations due to these infiltrating cells. To assess if both the presence and function of microglia regulated tissue mechanics, we employed the zebrafish mutant, Panther, a mutant which lacks a functional fms (M-CSF receptor) gene [43]. Colony stimulating factor 1 receptor (csf1ra) is a key determinant in microglia development[44–46]. In these fish, early macrophages differentiate and behave normally in the yolk sac[46], however, there is a delayed invasion of macrophages into the brain. Colonization of the larval brain is eventually achieved with ∼ 4-day delay[46]. We determined that the mutant showed a difference in brain morphology and gene expression at 4 dpf compared to WT (Figure 3a and b, Extended data 3a and b). Furthermore, there are decreased numbers of macrophages in the mutant brain compared to the WT (Figure 3a). Additionally, the blood brain barrier (BBB) is impaired in the mutant fish where increased leakage of 150kDa dextran is observed (Extended Data 3b) compared to WT (Extended Data 1a). OT measurements revealed that the regional heterogeneity observed in WT is lost where the magnitude of the complex modulus is comparable for all regions (Figure 3c and d). Direct comparison to WT revealed that the mutant brain is softer in all brain regions at 4 dpf (Figure 3e). Additionally, the fore- and hindbrain in the mutant become more liquid-like during development as the crossover frequency where viscous modulus (G’’) crosses over elastic modulus (G’) shifts to lower frequency (Figure 3c). G*, G’, and G” monotonically increase as a function of frequency and show power law dependence at frequencies >400Hz (Figure 3c, Extended Data 3c and d). However, the measured mechanical property of the midbrain showed modest increase during development concomitant with minimal changes in the exponent of the power law (Extended Data 3c-f). The normalized complex modulus data from whole frequency range by 3 dpf hindbrain revealed a more mechanically homogeneous brain (Figure 3d and e). A similar examination using Brillouin microscopy revealed that all brain regions at 4 dpf show reduced Brillouin shifts in the mutant compared to WT (Figure 3f). In addition to reduced macrophages, staining of acetylated tubulin was more homogenous over the development for panther mutant compared to WT development (Figure 2d, Figure 3g). We then performed similar calculations where we compared the raw and corrected correlation between Brillouin (longitudinal) modulus, M’ and OT elastic modulus G’ at 7Hz, 907 Hz and 15kHz in terms of the elastic modulus at development stage-4dpf for WT and mutant zebrafish (Figure 3h-i, Extended data 4). Comparison of linear fits between log(G’) for forebrain, midbrain, and hindbrain) and log(calculated Brillouin modulus M’) revealed poor correlation between techniques before corrections based on estimated brain refractive index and density (Figure 3h-i, Extended data 4). The overall correlations improved post correction for the measurements of 907Hz and 15kHz but no significant change for the measurements at 7Hz. Better fits were obtained for values measured at 7 and 907Hz compared to 15kHz (Figure 3i-j, Extended data 4). However, the fits were relatively low compared to the fits obtained post correction for the WT fish as a function of development (Figure 2h and i, Extended data 2).

**Figure 3:**
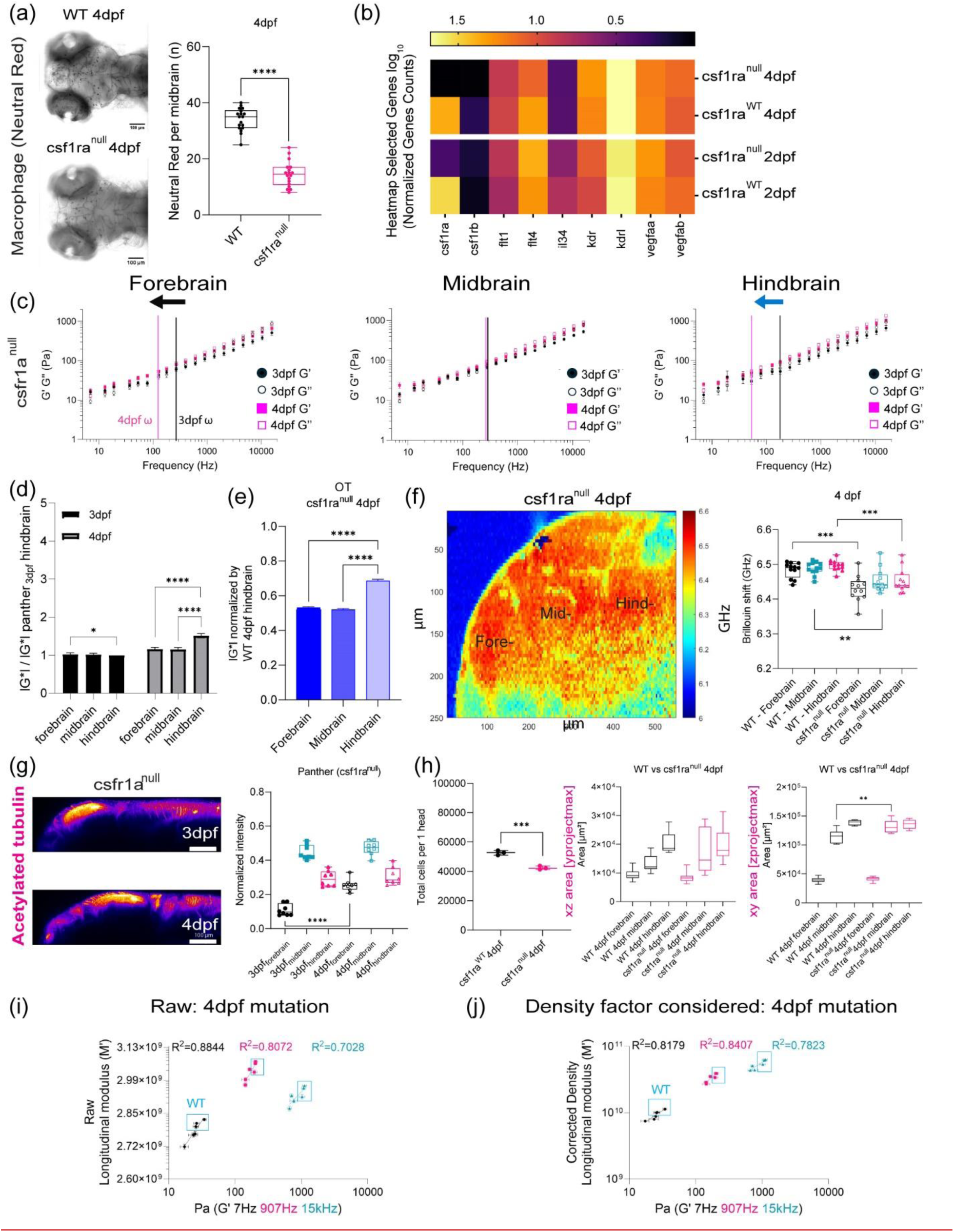
OT and Brillouin microscopy measured distinct mechanical differences in tissue mechanical properties in a zebrafish mutant characterized by impaired macrophage invasion. (a) Macrophages at midbrain labelled by neutral red assay comparing between WT and csf1ranull at 4dpf (WT 4dpf: n = 18 and csf1ra^null^ 4dpf: n = 18). **** p<0.0001, unpaired two-tailed t-tests (b) Bulk RNA sequence at 2dpf and 4dpf in terms of csf1ra mutation. (c) log-log plot of csf1ra^null^ brain mechanics (elastic modulus, G’, and viscous modulus, G’’) and frequencies (7Hz to 15kHz) in terms of development (3dpf and 4dpf) and brain regions (forebrain, midbrain, and hindbrain) with crossover frequency labelled where viscous modulus becomes greater than elastic modulus; 3dpf in black and 4dpf in magenta. Error bars in standard of error (3dpf = 4 fish – forebrain n (number of beads/fish) = 65, midbrain n = 88, and hindbrain n = 73. 4dpf = 5 fish – forebrain n (number of beads/fish) = 141, midbrain n = 140, and hindbrain n = 168). (d) normalized complex modulus, |G*|, of csf1ra^null^ at 19 different frequencies (7Hz to 15kHz) over the development in terms of brain regions normalized by 3dpf csf1ra hindbrain |G*|. (n=19), * p<0.05, **** p<0.0001, paired two-tailed t-test (e) normalized complex modulus, |G*|, of WT 4dpf at 19 different frequencies in respect to 3dpf WT hindbrain |G*| to show the degree of increase compared to csf1ra^null^. (n=19), **** p<0.0001, paired two-tailed t-test (f) 4dpf csf1ra^null^ Brillouin microscopy image with Brillouin shift (GHz) in terms of brain regions comparing with WT (WT = 12 and csf1ra^null^ = 12). ** p<0.01, ***p<0.001, unpaired two-tailed t-tests. (g) y-axis maximum projection of csf1ra^null^ acetylated tubulin over the development and its normalized intensity in terms of development and brain regions (3dpf n = 9 and 4dpf n = 7) (h) Dissociated head cells per fish based on the cluster of 15 fish in terms of csf1ra mutation at 4dpf (WT 4dpf: n = 45, csf1ra^null^ 4dpf: n = 45). Each dot represents a group of 15 fish head. *** p<0.001, unpaired two-tailed t-tests. Quantitation of brain region area based on acetylated tubulin in terms of csf1ra mutation at 4dpf, xz (y-axis maximum projection), and xy (z-axis maximum projection) (WT 4dpf n = 13 and csf1ra^null^ n = 7). ** p<0.01, unpaired two-tailed t-tests. (i) Raw correlation between corrected Brillouin (longitudinal) modulus (M’) and OT elastic modulus (G’) at 7Hz, 907Hz, and 15kHz at csf1ra mutation at 4dpf in log-log plot (j) Density factor corrected correlation between corrected Brillouin (longitudinal) modulus (M’) and OT elastic modulus (G’) at 7Hz, 907Hz, and 15kHz at csf1ra mutation at 4dpf in log-log plot.

### Acetylated tubulin is a major driver of brain mechanical properties

As immunofluorescence revealed that there is massive remodeling and increase in acetylated tubulin. After demonstrating that BM is sensitive to mechanical changes due to development and changes due to genetic mutants, we then investigated the role of acetylated tubulin in driving mechanical properties. Hence, we modulated expression of tubulin using two pharmacological inhibitors, Nocodozale and Blebbistatin. Nocodozale inhibits self-assembly of tubulin and/or associated proteins[47]. Moreover, it also depolymerizes preformed microtubules *in vivo* (Figure 4a-d). Brillouin frequency shifts decreased ∼ 150 – 400 MHz for all regions of the brain parenchyma compared to control (Figure 4a-b). We observed that the midbrain was more affected where we observed the greatest decrease in BM shift and acetylated tubulin upon treatment with nocodazole compared to the fore- and hindbrain (Figure 4 a-d). Myosin II plays a role in cell contractility which in turn can regulate tissue architecture. Furthermore, during development, Myosin II activity has been shown to contribute to spindle orientation for cell-division orientation in epiboly, a key developmental step for zebrafish[48]. Thus, we also probed the role of Myosin II in regulating the mechanical properties of the brain parenchyma using the pharmacological agent, Blebbistatin (Figure 4e-g). We confirmed that brain tissue was altered upon Blebbistatin treatment as acetylated tubulin staining showed a decreased normalized fluorescence intensity (Figure 4g). Inhibition of Myosin II activity revealed reduced Brillouin frequency shifts decreased about 15 – 50 MHz for all regions of the brain parenchyma compared to control (Figure 4e-f). Moreover, the regional trend in stiffness persisted as the hindbrain remains the stiffest region. However, the forebrain showed the largest decrease as we measured a Brillouin shift of 50 MHz reflecting the greatest observed reduction in stiffness. Acetylated tubulin intensity at the forebrain was most affected compared to the other regions. As we saw differences in macrophage infiltration as a function of development and mutant status, we then investigated if there would be difference upon inhibition of mechanical properties via modulation of tubulin. We determined that for each of these treatments there were differences in the number of macrophages that were present in the midbrain (Figure 4h).

**Figure 4:**
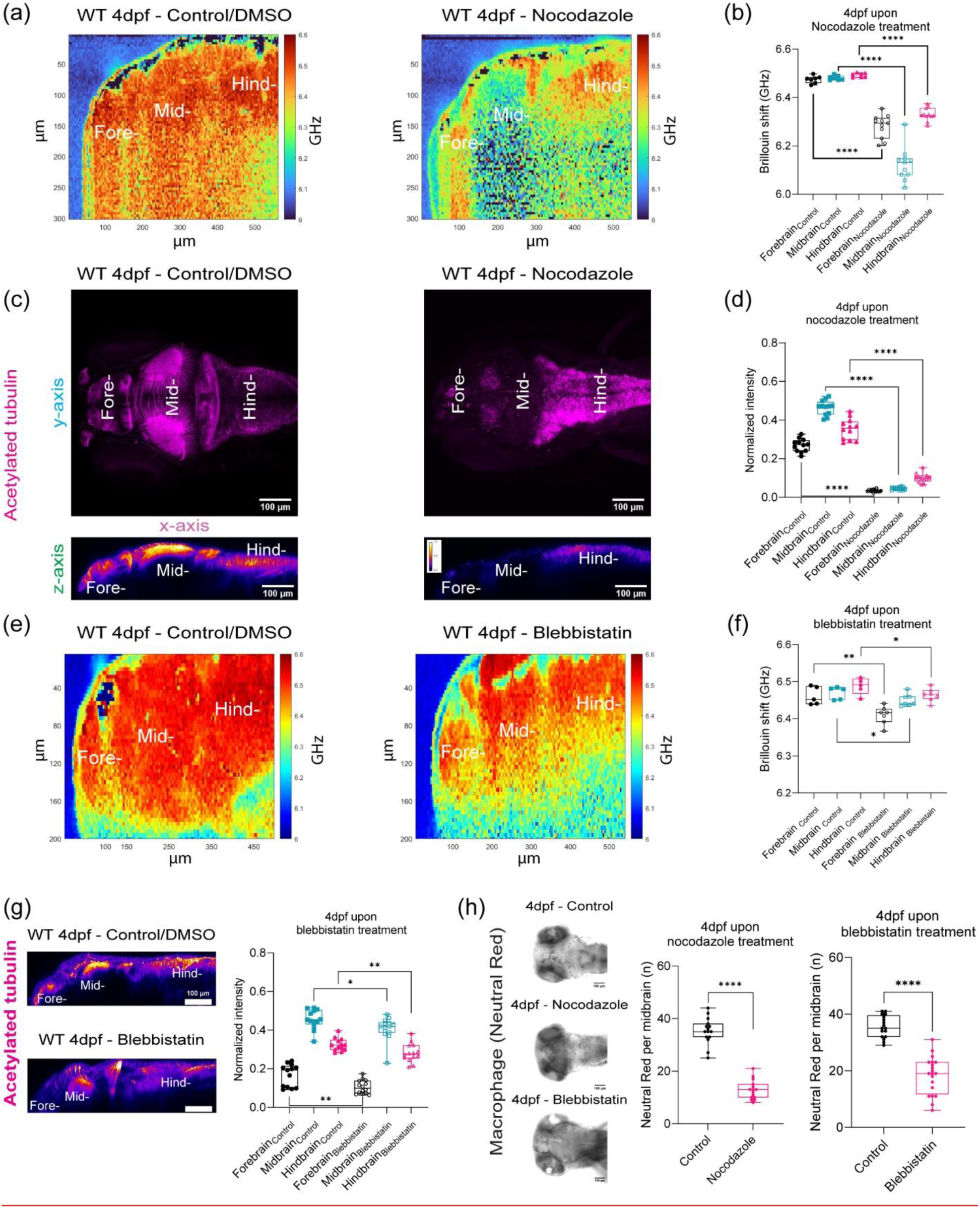
Acetylated tubulin is a major driver of brain mechanical properties. (a) Brillouin microscopy images for WT 4dpf control/DMSO and WT 4dpf Nocodazole treatment. (b) Brillouin shift comparison between WT 4dpf control/DMSO and WT 4dpf Nocodazole treatment in terms of brain regions. (control: n = 7 and Nocodazole: n = 11). **** p<0.0001, unpaired two tailed t-tests. (c) z-axis maximum projection (xy) and y-axis maximum projection (xz) for WT 4dpf acetylated tubulin in terms of Nocodazole treatment (d) Quantitation of normalized acetylated tubulin intensity in terms of brain regions upon Nocodazole treatment (control: n = 12 and Nocodazole: n = 13). **** p<0.0001, unpaired two-tailed t-tests. (e) Brillouin microscopy images for WT 4dpf control/DMSO and WT 4dpf Blebbistatin treatment (f) Brillouin shift comparison between WT 4dpf control/DMSO and WT 4dpf Blebbistatin treatment in terms of brain regions. (control: n = 5 and Blebbistatin: n = 7). * p<0.05, ** p<0.01, unpaired two tailed t-tests. (g) y-axis maximum projection (xz) of WT 4dpf acetylated tubulin in terms of Blebbistatin treatment and Quantitation of normalized acetylated tubulin intensity in terms of brain regions upon Blebbistatin treatment (control: n = 13 and Blebbistatin: n = 14). * p<0.05, ** p<0.01, unpaired two-tailed t-tests. (h) Macrophages at midbrain labelled by neutral red assay for WT upon Nocodazole treatment (WT 4dpf control: n = 15 and WT 4dpf Nocodazole: n = 16). **** p<0.0001, unpaired two-tailed t-tests. Macrophages at midbrain labelled by neutral red assay for WT upon Blebbistatin treatment (WT 4dpf control: n = 17 and WT 4dpf Blebbistatin: n = 20). **** p<0.0001, unpaired two-tailed t-tests.

### CSF1r macrophages density regulates brain mechanics

To confirm that the presence of macrophages or their remodeling capabilities can directly perturb mechanical properties of the brain, we ablated macrophages using two methods and blocked the receptor using pharmacological methods (Figure 5a-i). First, we injected nanoparticles (liposomes encapsulating clodronate) directly into the brain (Figure 5a-c). Second, we employed a chemogenetic model, Tg(mpeg1.1:NTR-IRES-eGFP-CAAX) where macrophages express an enhanced variant of Escherichia coli nitroreductase (NfsBT41Q/N71S/F124T) (NTR) resulting in macrophage specific ablation (Figure 5d-f) [49]. Upon treatment with Metronidazole, mpeg1.1^+^ cells are ablated. In each condition, we observed a reduction of macrophages. We then used a small molecule drug, PLX5622, against the csf1r receptor to assess if macrophage mediated tissue remodeling is important for mechanical homeostasis (Figure 5g-i). After 1 day post injection (dpi) of clodronate, there was ∼55% decrease in macrophage population and ∼ 65% reduction using the tissue specific ablation based on neutral red staining assay (Figure 5b and c) compared to control fish. Macrophage depletion due to clodronate liposomes, macrophage specific ablation, and CSF1R inhibitor (PLX5622) caused softening of whole brain parenchyma (Figure 5c, f, i) as all brain regions had a decrease in frequency shifts in the range of 70 – 90 MHz compared to control counterparts. The hindbrain showed the greatest reduction in stiffness concomitant with alterations in acetylated tubulin expression (Extended Data 5a-d). We observed a reduction of macrophages in the treated fish concomitant with a softening of all brain regions as measured using BM (Figure 5g-i). However, the greatest decrease in BM shift was observed for fish with the specific genetic ablation of the macrophages.

**Figure 5:**
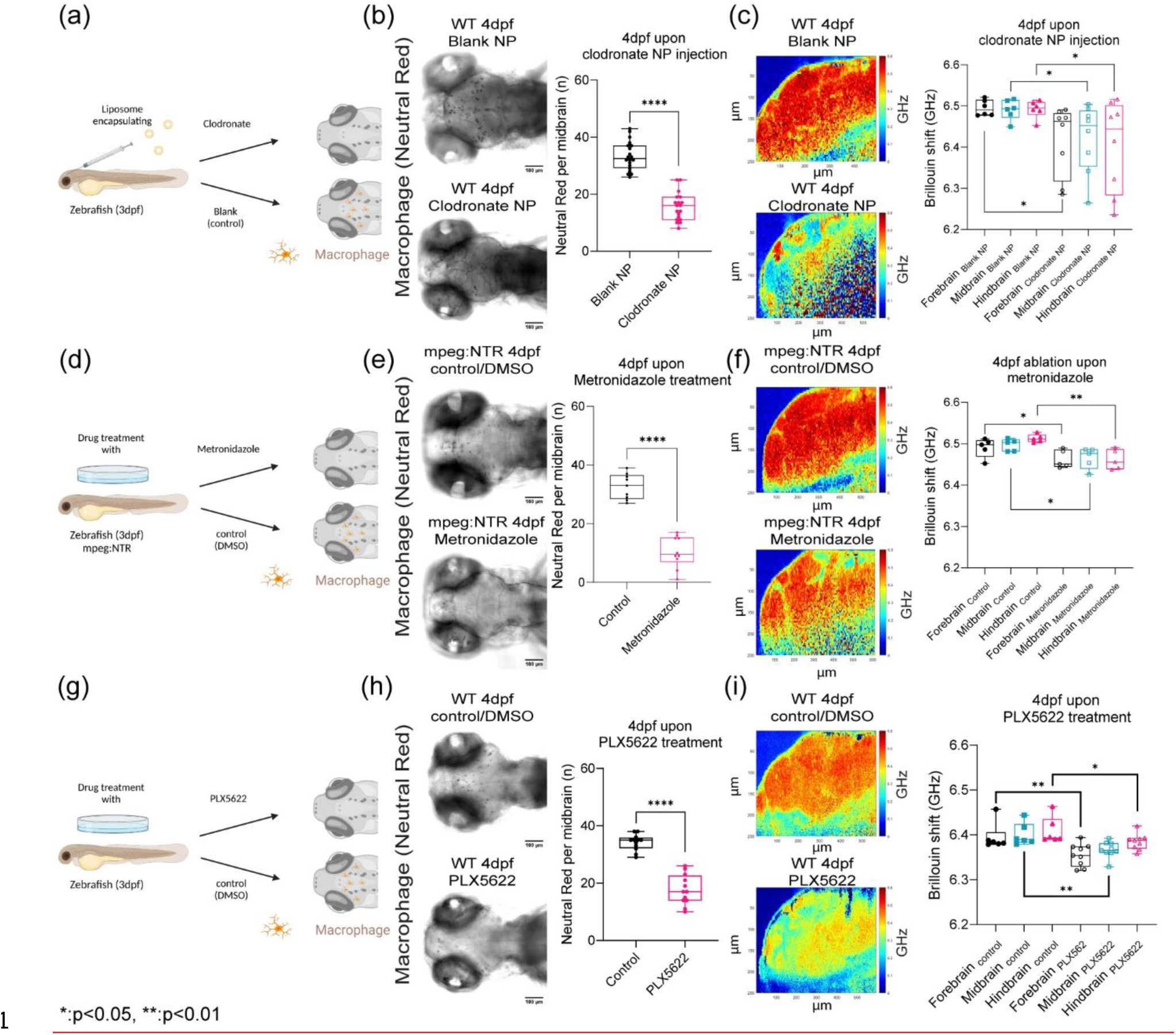
Macrophages and CSF1r-mediated stromal interactions regulate brain mechanics. (a) Macrophage depletion schematic by injecting nanoparticles (liposomes encapsulating clodronate) into WT 3dpf midbrain. (b) Macrophages at midbrain labelled by neutral red assay for WT 4dpf upon nanoparticle injection (WT 4dpf control NP: n = 20 and WT 4dpf clodronate NP: n = 19). **** p<0.0001, unpaired two-tailed t-tests (c) Brillouin shift comparison between WT 4dpf control NP and WT 4dpf clodronate NP in terms of brain regions. (control NP: n = 5 and clodronate NP: n = 8). * p<0.05, unpaired two tailed t-tests. (d) Macrophage depletion schematic by using mpeg:NTR fish upon Metronidazole treatment at 3dpf (e) Macrophages at midbrain labelled by neutral red assay at for mpeg:NTR 4dpf upon Metronidazole (control NP: n = 9 and WT 4dpf clodronate NP: n = 10). **** p<0.0001, unpaired two-tailed t-tests. (f) Brillouin shift comparison between 4dpf mpeg:NTR control/DMSO and 4dpf mpeg:NTR Metronidazole in terms of brain regions. (control: n = 5 and Metronidazole: n = 5). * p<0.05, ** p<0.01, unpaired two tailed t-tests. (g) Schematic of CSF1R inhibition using small molecule drug, PLX5622, on WT at 3dpf (h) Macrophages at midbrain labelled by neutral red assay at for WT 4dpf upon PLX5622 (control/DMSO: n = 12 and PLX5622: n = 12). **** p<0.0001, unpaired two-tailed t-tests. (i) Brillouin shift comparison between WT 4dpf control/DMSO and WT 4dpf PLX5622 in terms of brain regions. (control: n = 6 and PLX5622: n = 9). * p<0.05, ** p<0.01, unpaired two tailed t-tests.

## Discussion

Mechanical properties have been used as metrics to identify the presence of pathologies since antiquity where touch can sometimes serve as a sensile guide to assist in surgical resection of cancerous tissue[50]. Not restricted to cancers, a differential mechanical phenotype is also a hallmark of diseases such as fibrosis and diabetes[51]. Elastography methods, such as Brillouin scattering microscopy, form a class of non-invasive techniques that are used in preclinical models and in the clinic[21, 27, 52]. However, a key goal in using these techniques is establishing the mechanical signature obtained at the higher frequencies with our conventional understanding using other methods at frequency scales ∼kHz. Recent studies in the zebrafish show the broad utility of optical based techniques in probing biophysical properties such as viscoelasticity and hemodynamic forces *in vivo*[22, 29–31, 33, 40, 53, 54]. Here, we provide evidence for the biological origin of the BM scattering signal in the larval zebrafish brain. Our data suggest that these signals are largely driven by acetylated tubulin, microscale neuronal architecture, macrophage dynamics, and sub-cellular receptor modulation. Beyond linking BM to its biological determinants, we validated the extracted micro-mechanical signatures by performing a direct comparison against a gold-standard microscale mechanical test, OT-AMR. We first show that there is good agreement between the techniques despite the fact that they probe mechanical properties at different timescales. BM probes the longitudinal modulus on the GHz timescale which differs from both the Young’s and shear moduli measured by OT-AMR. Our results confirm congruence between traditional rheological methods in cells and tissues. There are some differences, such as the fact that a significant shift was not observed as the fish ages for BM measurements as observed for the OT-AMR (Figure 1d and f, Extended Data 1f). This could be due to the contributions of the “viscous” component of the material that changes as a function of aging as we observed regional differences in viscous modulus as a function of age, where the forebrain became more “liquid-like”, and the hindbrain became more “solid-like” from 3 to 4 dpf. In addition, the power law analysis revealed age dependent transitions between semi-flexible and flexible polymers at high frequencies (Extended Data 1c). One expression is that the cytoskeletal elements such as the tubulin change both stoichiometry and organization as a function of development. These results can possibly drive differences in the refractive indices and densities that are needed to interpret the longitudinal modulus as measured by BM (Figure 2h-i, Extended Data 2).

Recent advancements in contact-based atomic force microscopy (AFM) have generated valuable insights on tissue mechanics during embryogenesis[41, 55]. Our results confirm that morphogenetic programs such as invading immune cells and proliferation of neurons [18, 56–59]. Stromal cells and the associated signaling pathways associated with the sculpting embryonic tissue are also important in microenvironmental regulation of tumor etiology and metastatic colonization[17]. One such cell is the macrophage, a member of the innate immune system[17]. Here, we identified csfr1 macrophages as a key factor in brain tensional homeostasis. We determined that the mechanical properties are dependent on both the structural presence of the macrophages and the receptor that facilitates stromal remodeling. The mechanical homeostasis was modulated by both acute and permanent alterations of macrophage dynamics. Csf1r has pleiotropic functions in humans, rodents and zebrafish where loss of the receptor resulted in central nervous system and skeletal deformities[60]. In humans, variants contribute to a spectrum of development disorders that are classified collectively as BANDDOS: brain abnormalities, neurodegeneration and dysosteosclerosis[61]. More recently, variants of CSF1R have been implicated in adult-onset leukodystrophy[14]. The appearance of white matter lesions, axonal spheroids, and cerebral calcifications have deleterious effects on motor and cognitive functional resulting in death[14]. Although our results are obtained using larval fish, these observations may have consequences for adult tissues. An important area for future research exists in bridging the knowledge gap between our microscale techniques and the macroscale techniques such as magnetic resonance elastography which are already employed in the clinic.

Moreover, in cancers, tissue resident and recruited macrophages can help promote or suppress tumor growth, tumor cell escape, and distal organ colonization[16, 17]. Consequently, macrophage specific receptors have been pursued as druggable targets in the treatment of cancer [62]. These approaches have yielded mixed results in clinical studies. One missing factor could be that targeting the biochemical signaling of macrophages may alter mechanical properties of the microenvironment. These changes in turn could either act synergistically or blunt desired results of tumor clearance. Thus, it suggests that understanding the contribution of macrophage remodeling in the maintenance of mechanical homeostasis may be needed to harness macrophage-based therapeutics in disease management. Moreover, it underscores our need to understand normal tissue homeostasis if we are to design therapeutics that target the microenvironment. With this information in hand, we can begin to understand the functional role of tissue mechanics in the establishment and maintenance of normal tissue and what may go awry at the onset of and progression of disease.

## Methods

All animal experiments were done under protocols approved by the National Cancer Institute (NCI) and the National Institutes of Health (NIH) Animal Care and Use Committee.

### Zebrafish husbandry

Zebrafish were maintained at 28.5 °C on a 14-hour light/10-hour dark cycle according to standard procedures. Larvae were obtained from natural spawning, raised at 28.5 °C, and maintained in fish water, 60 mg sea salt (Instant Ocean, #SS15-10) per liter of DI water. For all experiments, larvae were transferred to fish water supplemented with N-phenylthiourea (PTU; Millipore Sigma, # P7629-25G) between 18-22 hours post-fertilization to inhibit melanin formation for enhanced optical transparency. PTU fish embryo water was prepared by dissolving 400 µl of PTU stock (7.5% w/v in DMSO) per 1 L of fish water. Water was replaced twice per day.

### Zebrafish lines

For wild type, AB strain zebrafish were crossed for brain mechanics measurement at 3 and 4 dpf. Additionally, csf1ra mutant zebrafish (Panther^j4as^) were crossed to validate the role of csf1ra on brain mechanics and development at 3 and 4 dpf. To investigate the role of macrophage on brain mechanics, Tg(mpeg1.1:NTR-IRES-eGFP-CAAX)^co57^ was crossed where macrophage, mpeg1.1 promoter, is expressed under eGFP. Tg(fli1a:eGFP)^y1^ zebrafish was crossed to visualize the integrity of blood-brain barrier upon 150kDa dextran circulation injection at 3 and 4 dpf compared to panther. For all experiments, at around 22 – 24 hpf, larvae were transferred to fish water supplemented with PTU to inhibit melanin formation for increased optical transparency. Larvae were then returned to the incubator at 28.5 °C and checked for normal development.

### Optical Tweezer (OT) based active microrheology measurement

Zebrafish larvae at 2 dpf were anesthetized using 0.4% buffered tricaine. 2nl of 1 µm polystyrene beads (ThermoFisher Scientific, #F8816) resuspended in PBS at a final concentration of 10^7^ beads/ml was injected directly into brain parenchyma (midbrain). Injected larvae were transferred into fresh PTU water and maintained at 28.5 °C. For OT measurement, injected larvae at 3 and 4dpf were anesthetized using 0.4% buffered tricaine and embedded in 1.25% low melting point agarose gel and allowed to polymerize in with cover glass (no. 1.5 thickness). Then, fish water supplemented with tricaine was added to the agarose hydrogel for the entire time of data acquisition as previously described[33, 34]. The details of our home-built OT setup are described[29, 40]. Mainly, the equipment is composed of two near infrared lasers, a 1064 nm trapping laser (IPG Photonics, #YLR 20 1064 Y11) to trap and oscillate a bead and a 975 nm detection laser (Lumics, #LU0975M00 1002F10D) to detect the movement of bead. Along with the two near infrared lasers, there are two quadrant photodiodes (QPD; First Sensor, #QP154-QHVSD) to detect trapping laser displacement (trap QPD) and the motion of bead (detection QPD). The trap laser is oscillated by a dual axis acousto-optic deflector (AOD; IntraAction, #DTD274HD6), which is operated by radio frequency generating cards (Analog Devices, #AD9854/PCBZ) with on-board temperature-controlled crystal oscillators[29, 40]. Then, the cards are digitally controlled by a data acquisition (DAQ) card (National Instruments, PCIe 5871R FPGA). Right after AOD, a small fraction of trap laser power (∼1%) is directed to trap QPD. Both trapping and detection lasers go through the backport of an inverted microscope (Nikon, Eclipse Ti-U) with a long working distance water immersive objective (Nikon, MRDO7602 CFIPLAN APO VC60XA WI 1.2 NA). Then, a long working distance (WD) and high numerical aperture (NA) condenser (Nikon, WI 0.9NA) collects the light from objective. After the condenser, a dichroic mirror (Chroma, ZT1064RDC-2P) reflects both trap and detection lasers. A bandpass filter (Chroma, #ET980/20x) excludes the trap laser so that the detection laser reaches detection QPD to record the motion of bead. Then, time-correlated trap and detection QPD signals from control and samples are collected and conducted by the DAQ card and custom program LabVIEW (National Instruments). During experiments, condenser is placed in Kohler illumination to locate beads via a piezo XYZ nanopositioning stage (Prior, #77011201) and a charge-coupled device (CCD) camera (Andor, Ixon DU-987E-C50-#BV). A selected bead is precisely positioned at the center of trap after scanning it by detection laser in three dimensions using a piezo nanopositioning while recording the voltages (V) from the detection QPD. The relationship between V and displacement (nm) relationship, β, from the detection QPD is calibrated *in situ* by fitting the central linear line of the detector responses to scanning the bead through the detection laser in the direction of the trap laser oscillations, yielding β in V/nm. A trap QPD records the position of the trap laser to find the relative phase lag between the trap oscillations and bead. The trap stiffness, *k*, is determined *in situ* for selected beads based on the active-passive calibration method [40]. As described by Fischer and Berg-Sørensen, the stiffness, *k*, is determined based on the active power spectrum, 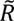_*L*_(ω), and passive power spectrum, *P*_*U*_(ω)[63]. The stiffness (Eq. 1) is

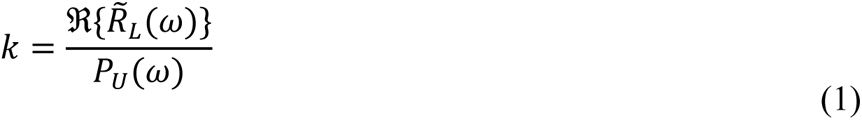

Active power spectrum, 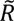_*L*_(ω), is recorded while trap laser is oscillating. The active power spectrum (Eq. 2) is

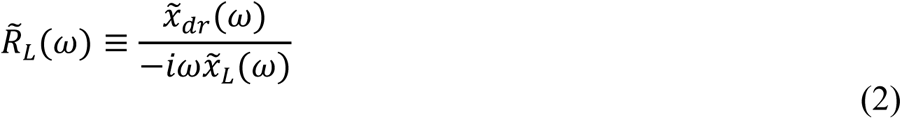

Where 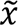_*L*_(ω) and 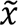_*dr*_(ω) are the Fourier transforms of the time series of the positions of the trap laser and the driven bead respectively.

Passive power spectrum, *P*_*U*_(ω), is measured while trap laser is held stationary. The passive power spectrum (Eq. 3) is

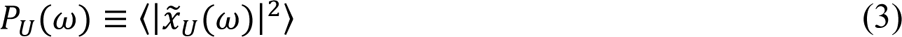

where 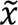_*U*_(ω) is the Fourier transform of the time series of the undriven bead’s thermally fluctuating position while trap laser is stationary.

With all the information from the trajectories of bead positions along with β, *k*, mass of bead (*m*), and radius (*a*), the generalized Stokes-Einstein relationship yield the complex modulus, *G* ∗ (ω), shown in (Eq. 4).

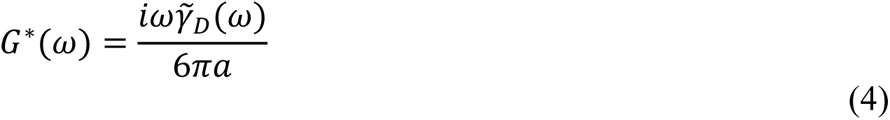

where 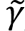_*D*_(ω) corresponds to the friction relaxation spectrum (Eq. 5) based on the active power spectrum, 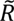_*L*_(ω).

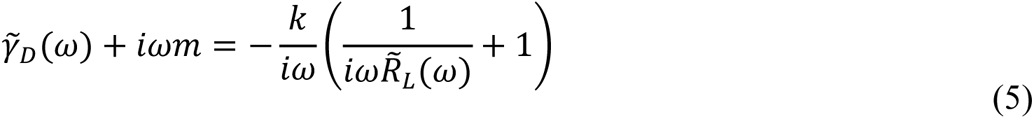

Then, the complex modulus can be broken into elastic modulus, *G*′(ω), and viscous modulus, *G*′′(ω) (Eq. 6).

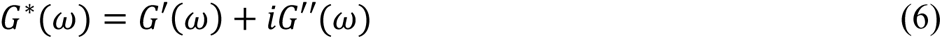

For all the measurements, laser power was set to 100 mW at the microscope back port while the amplitude of trap laser oscillations was set to 100nm. Experiments were controlled using custom LabVIEW programs.

### Brillouin microscopy

Larvae were anesthetized using 0.4% buffered tricaine and embedded in a dorsal orientation in 1.25% low melting point agarose gel and allowed to polymerize in with cover glass (no. 1.5 thickness) at 3 and 4 dpf. Dorsal orientation has been chosen to scan all brain regions (fore-, mid-, and hind-brain) but also to avoid eye interference during data acquisition. Then, fish water supplemented with tricaine was added to the agarose hydrogel for the entire time of measurement. Brillouin microscopy setup has been described elsewhere[23, 24]. Briefly, the setup is composed of 660 nm laser (Laser Quantum, #Torus-660), which illuminates the larvae brain at 20-30mW *via* 40x air objective (Olympus, LUCPLFLN40X 0.6 NA) after passing through the backport of microscopy body (Olympus, #IX81+DSU). Then, the backscattering from the focused voxel is collected through a single mode fiber (Thorlabs, #P1-460Y-FC-2), which serves as confocal pinhole. Brillouin light is then directed and analyzed by a two-stage VIPA etalon (FSR 15 GHz, LightMachinery, #OP-6721-6743-3) spectrometer in cross-axis configuration[19]. The VIPA spectrometer separates the frequencies of light, which is imaged on a high-sensitivity electron-multiplying charge-coupled device (EMCCD) camera (Andor, #iXon 897). Adjustable slits in the spectrometer blocks the stray light from elastic scattering to yield two Brillouin peaks, which are the anti-Stokes Brillouin scattering peak and the Stokes peak of the next diffraction order. At the end, the graph of intensity versus two Brillouin peaks (frequency) is obtained. The raw data is fitted with a Lorentzian function in a custom MATLAB program based on nonlinear least squares fitting to localize peak centers between the two Brillouin peaks. Brillouin microscopy measures spontaneous Brillouin scattering from the interactions between the incident light and inherent acoustic phonons (thermal fluctuations) inside the sample. Then, a frequency shift or Brillouin shift can be measured from the scattering. The Brillouin shift, *ν*_*B*_, is calculated based on (Eq. 7).

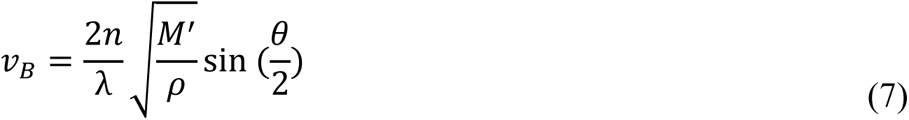

where *n* is refractive index of the material, *λ* is the laser wavelength, *M*′ is the longitudinal elastic modulus of measured sample, *ρ* is the density, and *θ* is the collection angle of the scattered light. For our setup, backward scattered light was collected, yielding *θ* = 180 °. Before actual sample measurements, Brillouin scattering calibration has been done by collecting 500 Brillouin spectra of methanol and water at 10ms exposure time. With the known literature values of Brillouin shift for methanol and water, the effective free spectral range and the spectral dispersion parameter (GHz per pixel) could be calculated. Calibration was done at least once an hour throughout whole measurements. The experimental parameters for brain scanning were done at 50-100ms exposure time and a pixel size at 2 µm x 2 µm.

Corrected Brillouin modulus was calculated based on the conversion of experimental Brillouin shifts from each brain regions to longitudinal modulus (*M*′). The longitudinal modulus is correlated to OT’s complex modulus (lG*l), elastic modulus (G’), and viscous modulus (G’’) at 7Hz, 907Hz, and 15kHz based on log-log linear relationship, log(*M*′) = a * log(lG*l, G’, or G’’) + b, where the relative change in Brillouin modulus is related to the relative change in OT G’. Thus, the slope (a) is multiplied to log(G’) in order to get corrected Brillouin modulus, where log(Brillouin modulus) = a * log(G’) + log(M’). During the conversion process, density (*ρ* = 1081 g/m^3^) and refractive index (*n* = 1.395) values[64] are kept constant to understand the correlation between OT and Brillouin.

### Drug treatments

- *Mysoin II inhibition*: (±)-Blebbistatin (Millipore Sigma, #203390-5MG) was diluted to 10 µM in 0.1% DMSO in PTU fish embryo water right before use to inhibit non muscle Myosin II. Larvae were treated overnight from 3 dpf. Thus, the control was 0.1% DMSO in PTU fish embryo water.

- *Acetylated tubulin perturbation*: Nocodazole (Selleckchem, #S2775) was diluted to 10 µM in 0.1% DMSO in PTU fish embryo water right before use to decrease acetylated tubulin. Larvae were treated overnight from 3 dpf. Thus, the control was 0.1% DMSO in PTU fish embryo water.

- *Macrophage ablation*: Metronidazole (Millipore Sigma, #M1547) was diluted to 10 mM in 0.1% DMSO in embryo medium immediately before use for nitroreductase-mediated cell ablation. The control was 0.1% DMSO in PTU fish embryo water. As Metronidazole is light sensitive, all treatment groups were kept in the incubator in the dark including control groups. The treatment started at 3dpf for overnight.

-*Macrophage depletion injection*: Zebrafish larvae at 3 dpf were anesthetized using 0.4% buffered tricaine. 1.2nl of clodronate liposomes (FormuMax, #F7010C-AH) was injected directly into brain parenchyma (midbrain) to deplete macrophages at the brain. As a control, 1.2nl of empty liposomes (FormuMax, #F70101-AH) was injected. Injected larvae were immediately transferred into fresh PTU fish embryo water.

-CSF1R inhibitor: PLX5622 (Selleckchem, #S8874-5mg) was diluted to 10 µM in 0.1% DMSO in PTU fish embryo water right before use to inhibit CSF1R. Larvae were treated overnight from 3 dpf. Thus, the control was 0.1% DMSO in PTU fish embryo water.

### Macrophage labeling

Neutral red (Millipore Sigma, # N7005) was diluted to 2.5 µg/ml in embryo medium and administered to 3-4 dpf larvae for 2.5 h at 28.5 °C in the dark. Following staining, larvae were washed two or three times in fresh PTU fish embryo water and monitored until nonspecific tissue redness had washed out (30 - 60 min), before mounting dorsal-down in low-melt agarose for brightfield imaging of optic tectum (midbrain) microglia.

### Zebrafish immunofluorescence and approximate brain density calculation based on brain area quantitation and the number of cells after dissociation

Larvae from 3 - 4dpf were fixed for 4 hours at 4°C in 4% paraformaldehyde in PBS. Fixed larvae were washed three times in PBDT (PBS supplemented with 1% DMSO and 0.5% Triton X-100). Washed larvae were gone sequential dehydration to methanol and then were stored at −20°C for at least 12°C hours. Dehydrated larvae in methanol were sequentially hydrated in PBDT. Then, larvae were permeabilized with 10 µg/ml Proteinase K (Millipore Sigma, #3115879001) in PBDT for 15 min at room temperature to remove epidermis, followed by 4% paraformaldehyde in PBS for 30 min. Larvae were washed three times with PBDT and blocked for 1 h at room temperature in PBDT containing 5% goat serum. Larvae were then incubated in a 1:200 dilution of mouse monoclonal anti-tubulin (acetal Lys40) antibody (GeneTex, #16292) in PBDT with 5% goat serum for at least three days at 4°C to stain acetylated tubulin. Stained larvae were washed quickly in PBDT three times and then washed an additional two times in PBDT for 15 min each. Antibody-stained larvae were then transferred to a 1:250 secondary antibody cocktail containing an AlexaFluor Plus 488 goat anti-mouse secondary antibody (Thermo Fisher Scientific, #A-11001) in PBDT with 5% goat serum. The larvae were stained for overnight at 4°C. Then, the fish were washed three times in PBDT for 15 min each and imaged with a Zeiss 780 LSM confocal microscopy.

Confocal z-stacks were acquired at 0.5 µm steps with the pinhole diameter set at 90.1 µm. 12-bit images were acquired with a Zeiss 20x EC Plan-Apochromat, 0.8 NA objective. Samples were excited with 488 nm light from an argon laser at 2% total power of 25 mW for normalized fluorescence intensity comparison. Transmittance spectrum was also recorded. The master gain was set at or below 650. Pixel dwell times of 1.58 ms were used. Then, images were max projected in terms of z-axis and y-axis based on Omer, et al. XYZ projection tool [65] for further image analysis by using ImageJ. Each brain region was quantified based on acetylated tubulin area to get volume change in terms of xz and xy area based on Marchant, et al. acetylated tubulin immunofluorescence normalization protocol[41]. The brain tissue was dissociated into single cells using Bresciani, et al. protocol for dissociation with 20ul Collagenase 100 mg/ml per 480ul 0.25% trypsin-EDTA per 15 zebrafish to get estimated brain tissue mass[66].

### High-Throughput Zebrafish Imaging via Vertebrate Automated Screening Technology (VAST)

At 2dpf, embryos were removed from the incubator Fish were put into flat-bottom 96-well plate in 150uL fish water containing PTU. Using LP Sampler (Union Biometrica), fish were loaded into capillary and imaged by a Vertebrate Automated Screening Technology (VAST) BioImager (Union Biometrica). The camera of the VAST BioImager was used to quantify brain development and morphology. The fish were scored based on development of the hindbrain. After acquisition, the embryos were dispensed into a 96-well plate, rinsed from tricaine, and placed back into the incubator.

### Bulk RNA sequencing

RNA from embryos was extracted using TriZol (Thermo Fisher Scientific Cat No 15596026) according to manufacturer protocol. 1.25ug of Total RNA was submitted for RNA-Sequencing to the NCI CCR Genomics Core. The libraries were made using Poly(A) Selection kit (illumina). 150bp fragments were sequenced by paired-end 75bp utilizing Illumina NextSeq. Reads were aligned to Ensembl Zebrafish Genome Build 11, GRCz11, using Histat21. SAMtools was used for sorting and indexing2. PartekFlow was used for ANOVA and pathway analysis.

### Statistical analysis

All data are displayed as representative or the results from multiple independent experiments. Data comparisons were made by paired two-tailed t-test to distinguish the heterogeneity of brain mechanics within brain regions and unpaired two-tailed t-test to determine the drug efficacy on brain mechanics in GraphPad Prism. Error bars are expressed as mean ± standard of error for optical tweezer data and as mean ± standard deviation for everything else, with * p < 0.05, ** p < 0.01, *** p < 0.001 and **** p < 0.0001.

## Acknowledgements

This effort was supported by the Intramural Research Program of the National Institutes of Health, the National Cancer Institute (KT). This work was also supported by grants from the National Science Foundation (DBI-1942003) and the National Institutes of Health (R21CA258008, R01HD095520) (GS). This work utilized the computational resources of the NIH HPC Biowulf cluster. (http://hpc.nih.gov) Services were provided by the CCR Genomics Core at National Cancer Institute. The panther zebrafish line and advice were kindly provided by Dr. David Parichy (University of Virginia). The Tg(mpeg1.1:NTR-IRES-eGFP-CAAX) was also kindly provided by Dr. Bruce Appel (University of Colorado, Denver.) We also thank Dr. Mayssa Mokalled (Washington University) St. Louis for helpful discussion. All schematics are created with BioRender.com.

---

**Extended Data1:**
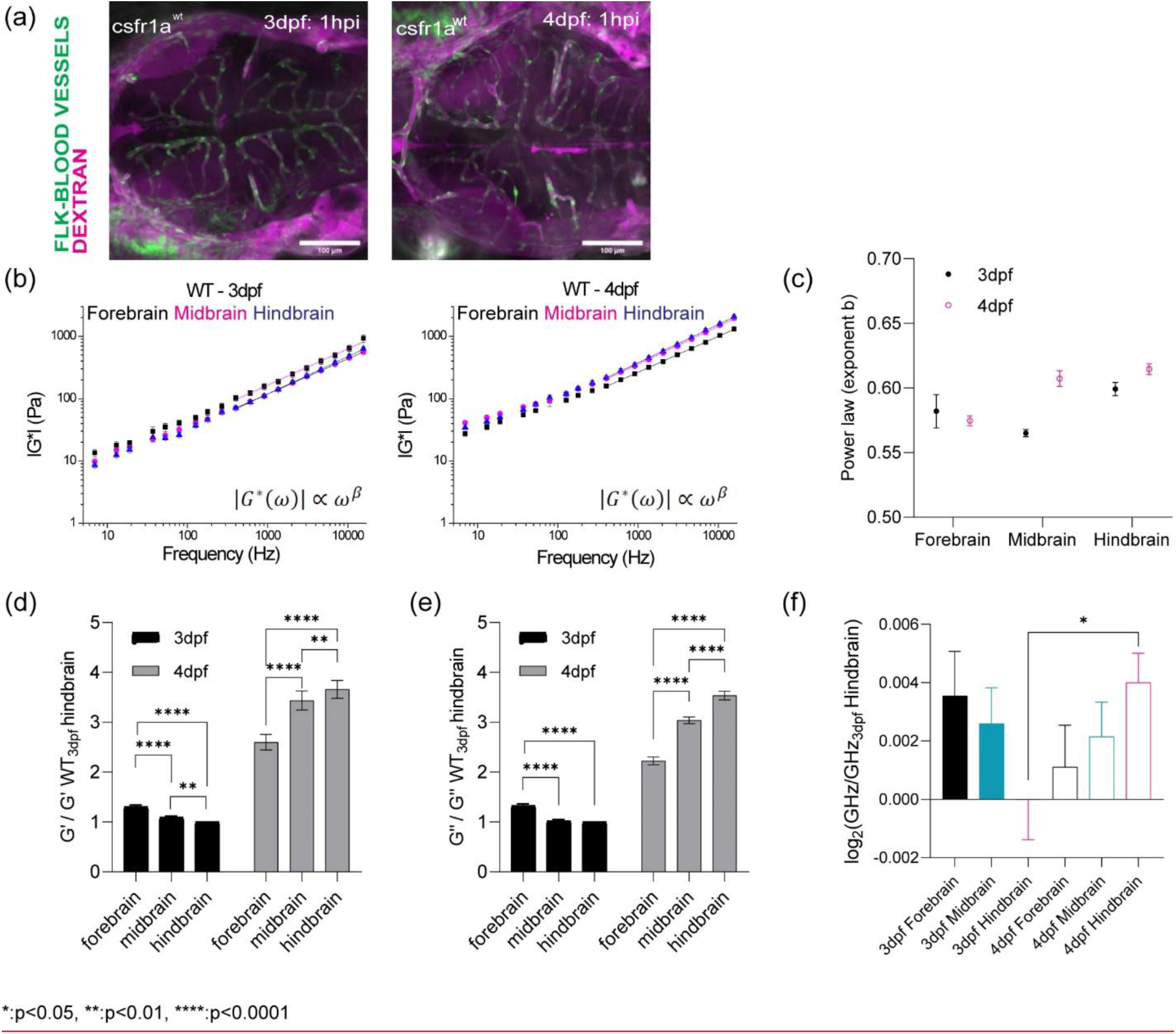
(a) Injection of 150kDa Dextran into blood circulation to look at the integrity of blood-brain barrier for WT in terms of development (b) log-log plot of complex modulus (G*) and frequencies from 7Hz to 15kHz with frequency-dependent power law fit (from 400Hz to 15kHz) (c) the exponent β of power law for WT in terms of development and brain regions (d) Normalized bar graph of WT G’ (elastic modulus) based on 19 different frequencies in respect to WT 3dpf hindbrain G’ (n = 19). ** p<0.01, **** p<0.0001, paired two-tailed t-tests. (e) Normalized bar graph of WT G’’ (viscous modulus) based on 19 different frequencies in respect to WT 3dpf hindbrain G’’ (n = 19). ** p<0.01, **** p<0.0001, paired two-tailed t-tests (f) log2 of normalized Brillouin shifts in respect to WT 3dpf hindbrain Brillouin shift (GHz) in terms of development and brain regions. * p<0.05, unpaired two-tailed t-tests.

**Extended Data2:**
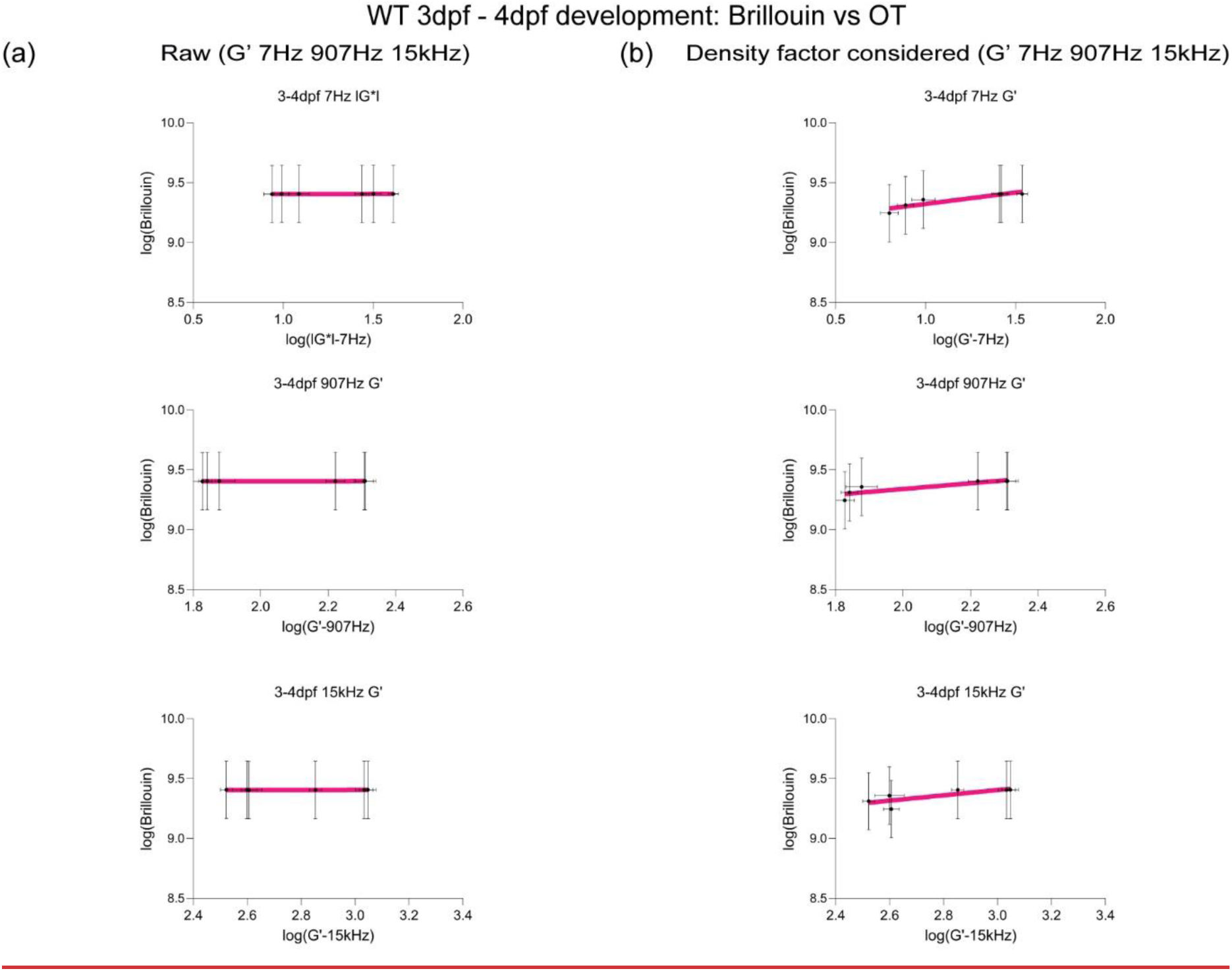
(a) Linear fit (top) between log(G’ for forebrain, midbrain, and hindbrain) and log(Brillouin modulus M’ calculated based on estimated brain refractive index and density) to get a slope to get corrected Brillouin modulus (bottom) in respect to G’ at 7Hz, 907Hz, and 15kHz during development (3-4dpf). (b) Linear fit (top) between log(G’ for forebrain, midbrain, and hindbrain) and log(Brillouin modulus M’ calculated based on estimated brain refractive index and density after density factor correction) to obtain corrected Brillouin modulus (bottom) in respect to G’ at 7Hz, 907Hz, and 15kHz during development (3-4dpf).

**Extended Data3:**
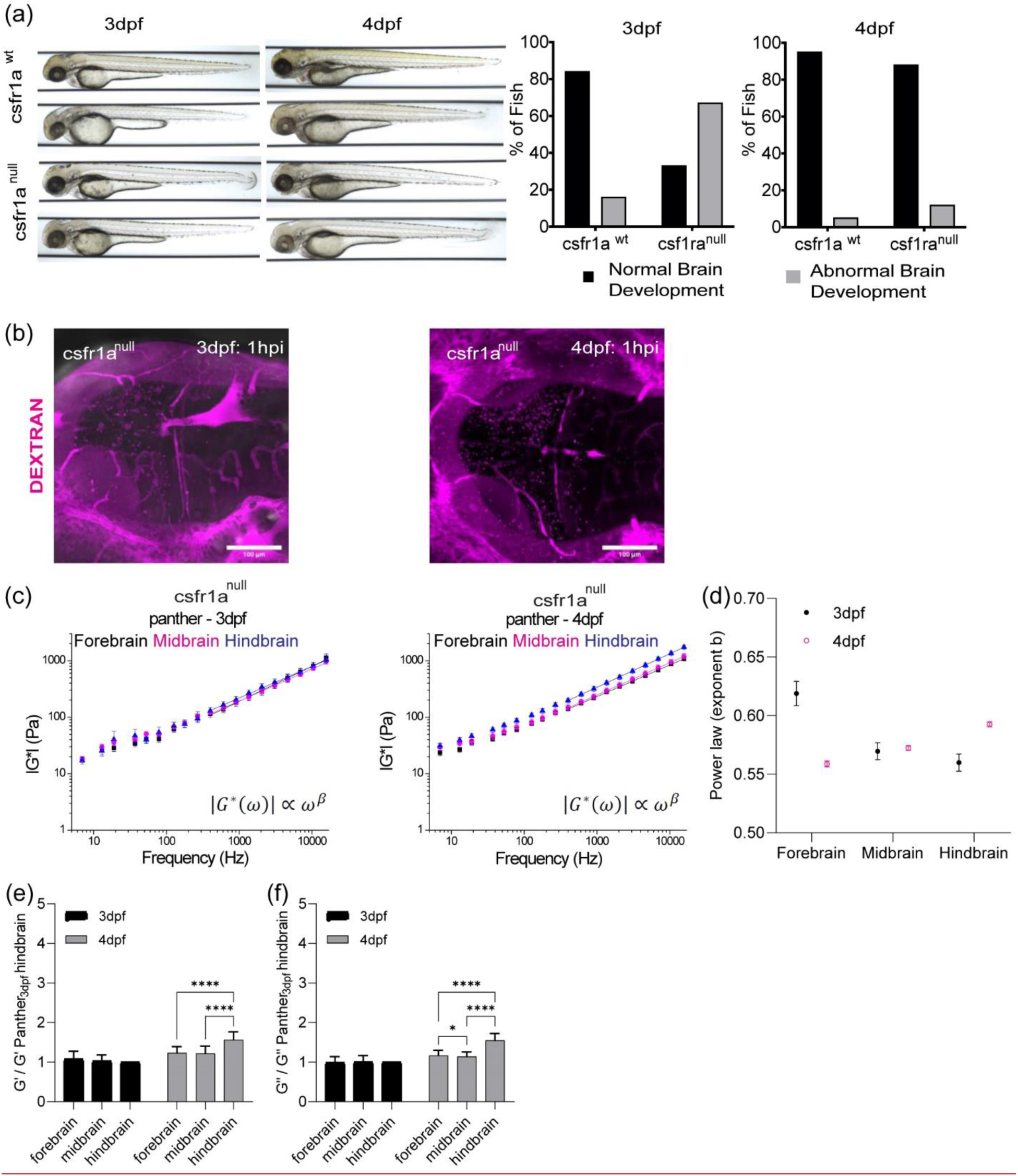
(a) The role of csf1ra on brain development at 3dpf and 4dpf (b) circulation injection of 150kDa Dextran to look at the integrity of blood-brain barrier for csf1ra^null^ in terms of development (c) log-log plot of csf1ra^null^ complex modulus (|G*|) and frequencies from 7Hz to 15kHz with frequency-dependent power law fit (from 400Hz to 15kHz) (d) the exponent β of power law for WT in terms of development and brain regions. (e) normalized bar graph of csf1ra^null^ G’ (elastic modulus) based on 19 different frequencies in respect to csf1ra^null^ 3dpf hindbrain G’ (n = 19). **** p<0.0001, paired two-tailed t-tests. (f) Normalized bar graph of csf1ra^null^ G’’ (viscous modulus) based on 19 different frequencies in respect to csf1ra^null^ 3dpf hindbrain G’’ (n = 19). * p<0.05, **** p<0.0001, paired two-tailed t-tests.

**Extended Data4:**
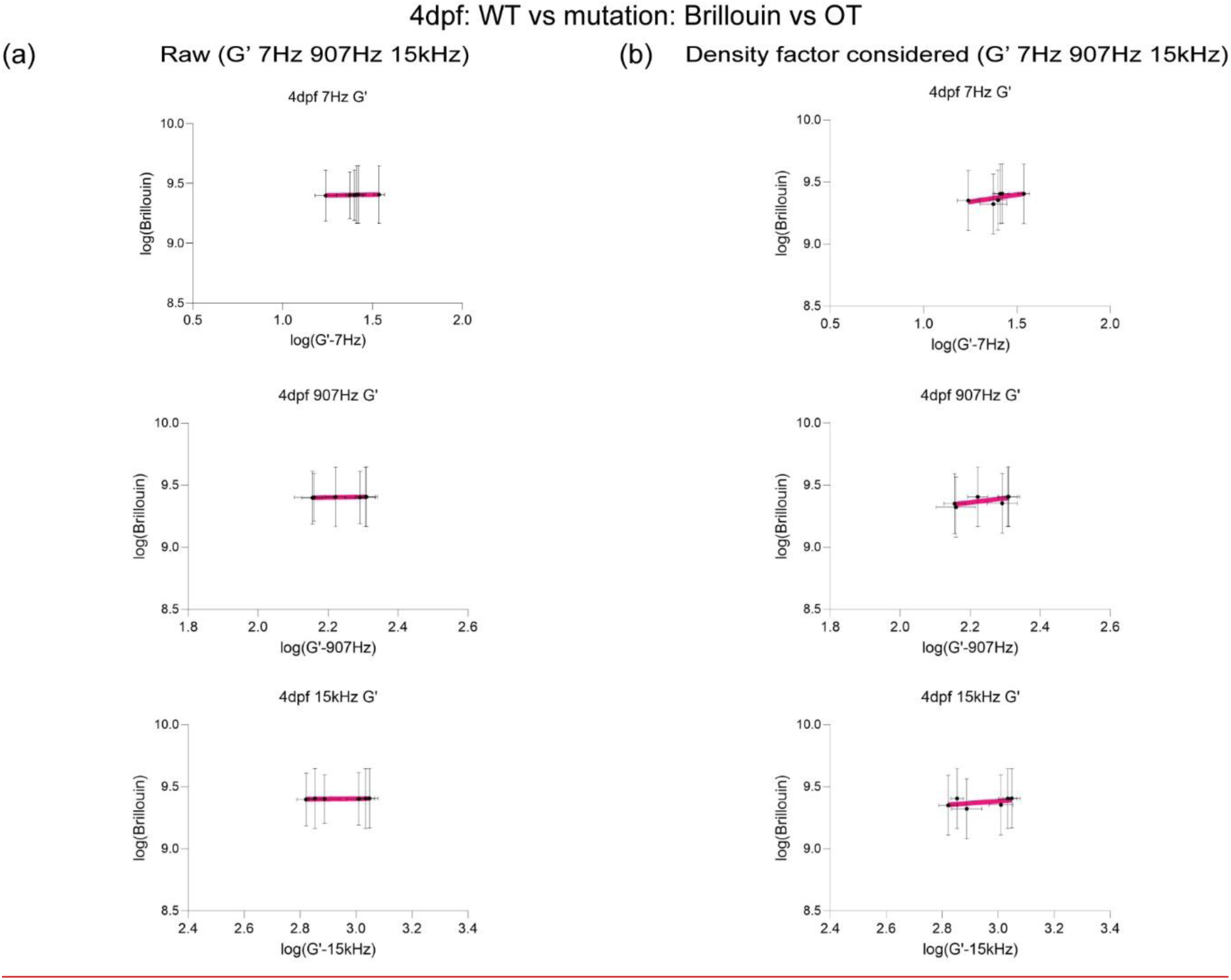
(a) Linear fit (top) between log(G’ for forebrain, midbrain, and hindbrain) and log(Brillouin modulus M’ calculated based on estimated brain refractive index and density) to obtain corrected Brillouin modulus (bottom) in respect to G’ at 7Hz, 907Hz, and 15kHz for csf1ra mutation at 4dpf. (b) Linear fit (top) between log(G’ for forebrain, midbrain, and hindbrain) and log(Brillouin modulus M’ calculated based on estimated brain refractive index and density after density factor correction) to get a slope to get corrected Brillouin modulus (bottom) in respect to G’ at 7Hz, 907Hz, and 15kHz for csf1ra mutation at 4dpf.

**Extended Data5:**
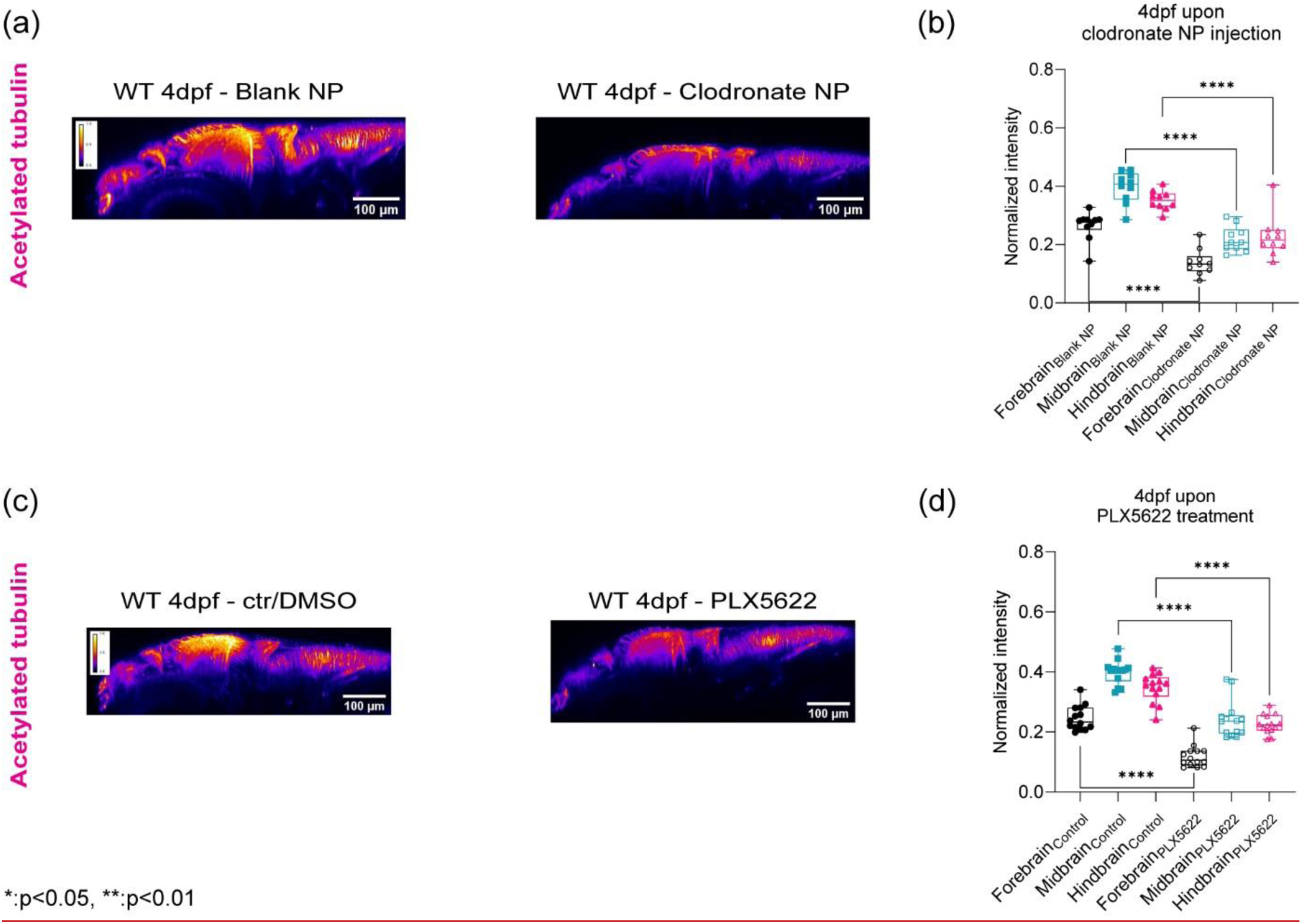
(a) y-axis maximum projection (xz) of WT 4dpf acetylated tubulin in terms of clodronate NP injection (b) Quantitation of normalized acetylated tubulin intensity in terms of brain regions upon clodronate NP injection (control NP: n = 12 and clodronate NP: n = 10). **** p<0.0001, unpaired two-tailed t-tests. (c) y-axis maximum projection (xz) of WT 4dpf acetylated tubulin in terms of control/DMSO and PLX5622 (b) Quantitation of normalized acetylated tubulin intensity in terms of brain regions upon PLX5622 treatment (control/DMSO: n = 14 and PLX5622: n = 13). **** p<0.0001, unpaired two-tailed t-tests.

## References

1. Barnes, J.M., L. Przybyla, and V.M. Weaver, Tissue mechanics regulate brain development, homeostasis and disease. J Cell Sci, 2017. 130(1): p. 71–82.

2. Keung, A.J., K.E. Healy, S. Kumar, and D.V. Schaffer, Biophysics and dynamics of natural and engineered stem cell microenvironments. Wiley Interdiscip Rev Syst Biol Med, 2010. 2(1): p. 49–64.

3. Klatt, D., et al., Noninvasive assessment of the rheological behavior of human organs using multifrequency MR elastography: a study of brain and liver viscoelasticity. Phys Med Biol, 2007. 52(24): p. 7281–94.

4. Budday, S., et al., Mechanical properties of gray and white matter brain tissue by indentation. J Mech Behav Biomed Mater, 2015. 46: p. 318–30.

5. Chaudhuri, O., et al., Effects of extracellular matrix viscoelasticity on cellular behaviour. Nature, 2020. 584(7822): p. 535–546.

6. Kasza, K.E., et al., The cell as a material. Current Opinion in Cell Biology, 2007. 19(1): p. 101–107.

7. Nobs, S.P. and M. Kopf, Tissue-resident macrophages: guardians of organ homeostasis. Trends in Immunology, 2021. 42(6): p. 495–507.

8. Yu, F.-X., B. Zhao, and K.-L. Guan, Hippo Pathway in Organ Size Control, Tissue Homeostasis, and Cancer. Cell, 2015. 163(4): p. 811–828.

9. Kiecker, C. and A. Lumsden, Compartments and their boundaries in vertebrate brain development. Nature Reviews Neuroscience, 2005. 6(7): p. 553–564.

10. Daneman, R. and A. Prat, The blood-brain barrier. Cold Spring Harb Perspect Biol, 2015. 7(1): p. a020412.

11. Schafer, D.P. and B. Stevens, Microglia Function in Central Nervous System Development and Plasticity. Cold Spring Harb Perspect Biol, 2015. 7(10): p. a020545.

12. Louveau, A., et al., Understanding the functions and relationships of the glymphatic system and meningeal lymphatics. J Clin Invest, 2017. 127(9): p. 3210–3219.

13. Cserép, C., et al., Microglia monitor and protect neuronal function through specialized somatic purinergic junctions. Science, 2020. 367(6477): p. 528–537.

14. Berdowski, W.M., et al., Dominant-acting CSF1R variants cause microglial depletion and altered astrocytic phenotype in zebrafish and adult-onset leukodystrophy. Acta Neuropathol, 2022. 144(2): p. 211–239.

15. Mattiace, L.A., P. Davies, S.H. Yen, and D.W. Dickson, Microglia in cerebellar plaques in Alzheimer’s disease. Acta Neuropathol, 1990. 80(5): p. 493–8.

16. Cassetta, L. and J.W. Pollard, Targeting macrophages: therapeutic approaches in cancer. Nat Rev Drug Discov, 2018. 17(12): p. 887–904.

17. Ruffell, B. and L.M. Coussens, Macrophages and therapeutic resistance in cancer. Cancer Cell, 2015. 27(4): p. 462–72.

18. Thompson, D.W., On growth and form. 1942: Cambridge Univ. Press. 116 pp.

19. Scarcelli, G. and S.H. Yun, *Multistage VIPA, etal*ons for high-extinction parallel Brillouin spectroscopy. Optics Express, 2011. 19(11): p. 10913–10922.

20. Scarcelli, G., P. Kim, and Seok H. Yun, In Vivo Measurement of Age-Related Stiffening in the Crystalline Lens by Brillouin Optical Microscopy. Biophysical Journal, 2011. 101(6): p. 1539–1545.

21. Scarcelli, G., R. Pineda, and S.H. Yun, Brillouin optical microscopy for corneal biomechanics. Invest Ophthalmol Vis Sci, 2012. 53(1): p. 185–90.

22. Schlussler, R., et al., Mechanical Mapping of Spinal Cord Growth and Repair in Living Zebrafish Larvae by Brillouin Imaging. Biophys J, 2018. 115(5): p. 911–923.

23. Nikolic, M. and G. Scarcelli, Long-term Brillouin imaging of live cells with reduced absorption-mediated damage at 660 nm wavelength. Biomed Opt Express, 2019. 10(4): p. 1567–1580.

24. Zhang, J. and G. Scarcelli, Mapping mechanical properties of biological materials via an add-on Brillouin module to confocal microscopes. Nat Protoc, 2021. 16(2): p. 1251–1275.

25. Nikolic, M., G. Scarcelli, and K. Tanner, Multimodal microscale mechanical mapping of cancer cells in complex microenvironments. Biophys J, 2022. 121(19): p. 3586–3599.

26. Bevilacqua, C., et al., High-resolution line-scan Brillouin microscopy for live imaging of mechanical properties during embryo development. Nat Methods, 2023.

27. Zhang, H., et al., Motion Tracking Brillouin Microscopy Evaluation of Normal, Keratoconic, and Post-Laser Vision Correction Corneas: Motion Tracking Brillouin Microscopy in Keratoconus and Laser Vision Correction. American Journal of Ophthalmology, 2023.

28. Zhang, J., M. Nikolic, K. Tanner, and G. Scarcelli, Publisher Correction: Rapid biomechanical imaging at low irradiation level via dual line-scanning Brillouin microscopy. Nat Methods, 2023.

29. Blehm, B.H., A. Devine, J.R. Staunton, and K. Tanner, In vivo tissue has non-linear rheological behavior distinct from 3D biomimetic hydrogels, as determined by AMOTIV microscopy. Biomaterials, 2016. 83: p. 66–78.

30. Harlepp, S., F. Thalmann, G. Follain, and J.G. Goetz, Hemodynamic forces can be accurately measured in vivo with optical tweezers. Mol Biol Cell, 2017. 28(23): p. 3252–3260.

31. Johansen, P.L., et al., Optical micromanipulation of nanoparticles and cells inside living zebrafish. Nat Commun, 2016. 7: p. 10974.

32. Lieschke, G.J. and P.D. Currie, Animal models of human disease: zebrafish swim into view. Nat Rev Genet, 2007. 8(5): p. 353–67.

33. Paul, C.D., et al., Tissue Architectural Cues Drive Organ Targeting of Tumor Cells in Zebrafish. Cell Syst, 2019. 9(2): p. 187–206 e16.

34. Paul, C.D., et al., Human macrophages survive and adopt activated genotypes in living zebrafish. Sci Rep, 2019. 9(1): p. 1759.

35. White, R., K. Rose, and L. Zon, Zebrafish cancer: the state of the art and the path forward. Nat Rev Cancer, 2013. 13(9): p. 624–36.

36. Renshaw, S.A. and N.S. Trede, A model 450 million years in the making: zebrafish and vertebrate immunity. Dis Model Mech, 2012. 5(1): p. 38–47.

37. Fabry, B., et al., Scaling the microrheology of living cells. Phys Rev Lett, 2001. 87(14): p. 148102.

38. Trepat, X., G. Lenormand, and J.J. Fredberg, Universality in cell mechanics. Soft Matter, 2008. 4(9): p. 1750–1759.

39. Staunton, J.R., W.Y. So, C.D. Paul, and K. Tanner, High-frequency microrheology in 3D reveals mismatch between cytoskeletal and extracellular matrix mechanics. Proc Natl Acad Sci U S A, 2019. 116(29): p. 14448–14455.

40. Staunton, J.R., B. Blehm, A. Devine, and K. Tanner, In situ calibration of position detection in an optical trap for active microrheology in viscous materials. Opt Express, 2017. 25(3): p. 1746–1761.

41. Marchant, C.L., A.N. Malmi-Kakkada, J.A. Espina, and E.H. Barriga, Cell clusters softening triggers collective cell migration in vivo. Nature Materials, 2022. 21(11): p. 1314–1323.

42. Chitnis, A. and J. Kuwada, Axonogenesis in the brain of zebrafish embryos. The Journal of Neuroscience, 1990. 10(6): p. 1892–1905.

43. Parichy, D.M., et al., An orthologue of the kit-related gene fms is required for development of neural crest-derived xanthophores and a subpopulation of adult melanocytes in the zebrafish, Danio rerio. Development, 2000. 127(14): p. 3031–3044.

44. Li, Q. and B.A. Barres, Microglia and macrophages in brain homeostasis and disease. Nat Rev Immunol, 2018. 18(4): p. 225–242.

45. Ferrero, G., M. Miserocchi, E. Di Ruggiero, and V. Wittamer, A csf1rb mutation uncouples two waves of microglia development in zebrafish. Development, 2021. 148(1).

46. Herbomel, P., B. Thisse, and C. Thisse, Zebrafish Early Macrophages Colonize Cephalic Mesenchyme and Developing Brain, Retina, and Epidermis through a M-CSF Receptor-Dependent Invasive Process. Developmental Biology, 2001. 238(2): p. 274–288.

47. Xie, R., S. Nguyen, W.L. McKeehan, and L. Liu, Acetylated microtubules are required for fusion of autophagosomes with lysosomes. BMC Cell Biology, 2010. 11(1): p. 89.

48. Campinho, P., et al., Tension-oriented cell divisions limit anisotropic tissue tension in epithelial spreading during zebrafish epiboly. Nat Cell Biol, 2013. 15(12): p. 1405–14.

49. Hughes, A.N. and B. Appel, Microglia phagocytose myelin sheaths to modify developmental myelination. Nature Neuroscience, 2020. 23(9): p. 1055–1066.

50. Michor, F., J. Liphardt, M. Ferrari, and J. Widom, What does physics have to do with cancer? Nat Rev Cancer, 2011. 11(9): p. 657–70.

51. So, W.Y. and K. Tanner, Emerging principles of cancer biophysics. Fac Rev, 2021. 10: p. 61.

52. Jordan, J.E.L., et al., Microscopic multifrequency MR elastography for mapping viscoelasticity in zebrafish. Magn Reson Med, 2022. 87(3): p. 1435–1445.

53. Bambardekar, K., et al., Direct laser manipulation reveals the mechanics of cell contacts in vivo. Proc Natl Acad Sci U S A, 2015. 112(5): p. 1416–21.

54. Serwane, F., et al., In vivo quantification of spatially varying mechanical properties in developing tissues. Nat Methods, 2017. 14(2): p. 181–186.

55. Thompson, A.J., et al., Rapid changes in tissue mechanics regulate cell behaviour in the developing embryonic brain. eLife, 2019. 8: p. e39356.

56. Lu, P., K. Takai, V.M. Weaver, and Z. Werb, Extracellular matrix degradation and remodeling in development and disease. Cold Spring Harb Perspect Biol, 2011. 3(12).

57. Petridou, N.I., Z. Spiro, and C.P. Heisenberg, Multiscale force sensing in development. Nat Cell Biol, 2017. 19(6): p. 581–588.

58. Shook, D. and R. Keller, Mechanisms, mechanics and function of epithelial-mesenchymal transitions in early development. Mech Dev, 2003. 120(11): p. 1351–83.

59. Vu, T.H. and Z. Werb, Matrix metalloproteinases: effectors of development and normal physiology. Genes Dev, 2000. 14(17): p. 2123–33.

60. Hume, D.A., et al., Phenotypic impacts of CSF1R deficiencies in humans and model organisms. Journal of Leukocyte Biology, 2019. 107(2): p. 205–219.

61. Guo, L. and S. Ikegawa, From HDLS to BANDDOS: fast-expanding phenotypic spectrum of disorders caused by mutations in CSF1R. Journal of Human Genetics, 2021. 66(12): p. 1139–1144.

62. Cannarile, M.A., et al., Colony-stimulating factor 1 receptor (CSF1R) inhibitors in cancer therapy. Journal for ImmunoTherapy of Cancer, 2017. 5(1): p. 53.

63. Fischer, M. and K. Berg-Sørensen, Calibration of trapping force and response function of optical tweezers in viscoelastic media. Journal of Optics A: Pure and Applied Optics, 2007. 9(8): p. S239.

64. Tapan K Viswas, T.M.L., *In vivo MR Measurement of Refractive Index, Relative Water Content and T2 Relaxation time of Various Brain lesions With Clinical Application to Discriminate Brain Lesions*. The Internet Journa of Radiology, 2009. 13(1): p. 1–9.

65. Omer, S., S.R. Greenberg, and W.-L. Lee, Cortical dynein pulling mechanism is regulated by differentially targeted attachment molecule Num1. eLife, 2018. 7: p. e36745.

66. Bresciani, E., E. Broadbridge, and P.P. Liu, An efficient dissociation protocol for generation of single cell suspension from zebrafish embryos and larvae. MethodsX, 2018. 5: p. 1287–1290.

